# Conservation implications of diverse demographic histories: the case study of green peafowl (*Pavo muticus*, Linnaeus 1766)

**DOI:** 10.1101/2023.07.21.549982

**Authors:** Ajinkya Bharatraj Patil, Nagarjun Vijay

**Affiliations:** Computational Evolutionary Genomics Lab, Department of Biological Sciences, IISER Bhopal, Bhauri, Madhya Pradesh, India

**Author notes:** **Name and complete mailing address:** Nagarjun Vijay Academic Building III (CELL) Indian Institute of Science Education and Research Bhopal Indore By-pass Road Bhauri Bhopal, 462066 Madhya Pradesh, India.

**Keywords:** Conservation, demographic history, endangered species, Evolutionarily Significant Units (ESUs), green peafowl, population structure

## Abstract

The green peafowl (*Pavo muticus*, Linnaeus 1766) is an endangered species native to the forests of tropical Southeast Asia. Although its morphological diversity and subspecies categorization is known and built upon traditional taxonomy, the intraspecific genetic structure has not been comprehensively addressed. To assess if phenotypic diversity is reflected at the molecular level, we used public whole-genome sequencing data of one blue peafowl and 52 green peafowls from multiple countries to characterize their genetic diversity, differentiation, identify Ancestry Informative Markers (AIMs) and compare their demographic histories. We found evidence of substantial population structure, with at least three distinct clusters and diverse demographic histories that may mirror different responses to various biogeoclimatic events. The genetic structure of native populations follows the pattern of the geographic distribution of the green peafowl with the highest autosomal pairwise F_ST_ between Yunnan and Vietnam (∼0.1) and intermediate estimates for Thailand comparisons (∼0.077). We identify AIMs to distinguish between these three native populations. The captive green peafowls from Xinxing clustered with Vietnam and those from Qinhuangdao (QHD) formed a separate cluster. The two QHD individuals appear to have varying levels of blue peafowl ancestry based on PCA and admixture analysis and are mirrored in their demographic histories. Our study establishes the occurrence of genetically distinct natural populations of green peafowl that can be considered separate management units (MU) when planning conservation actions. Transboundary cooperation and concerted efforts to foster genetic diversity are imperative for Southeast Asian species at risk.

## Introduction

The ongoing rapid loss of biodiversity due to the anthropogenic degradation of ecosystems, especially in tropical developing economies, is a major global issue with socioeconomic implications (Mikkelson et al. 2007; Johnson et al. 2017). While Southeast Asia hosts several biodiversity hotspots, it also has one of the highest deforestation rates, which could be a potential cause of massive ongoing and future species loss (Sodhi et al. 2004). The conservation of biodiversity necessitates the preservation of the natural habitat, which frequently conflicts with the objectives of economic development and presents a dilemma for the stakeholders (Sih et al. 2000). One of the prominent examples of habitat-loss mediated species decline across its distribution due to anthropogenic changes is the green peafowl (van Balen et al. 1995; McGowan et al. 1998; Brickle 2002; Nuttall et al. 2017; Sukumal et al. 2017, 2020; Kong et al. 2018; Saridnirun et al. 2021). Due to the rapid decline of this species and localized extinction within its native range, it has been classified as endangered, and conservation efforts are ongoing.

Historically, the green peafowl (*Pavo muticus*, Linnaeus 1766) was distributed across Southeast Asia, ranging from Northeast India to the island of Java (McGowan et al. 1998). At least three well-defined subspecies are known based on their morphology and distribution in the Javan (*P. m. muticus*), Indo-Chinese (*P. m. imperator*), and Burman (*P. m. spicifer*) regions (Delacour et al. 1977). The sub-species *P. m. spicifer* has been described from Myanmar, with a distribution extending to northeast India. The current distribution of this sub-species needs to be assessed in detail. The *P. m. muticus* was historically distributed in Malay peninsula and Java. It is now considered extinct in Malaysia as assessed by IUCN, whereas it has a surviving population in Java (Hernowo et al. 2011). The *P. m. imperator* appears to be the most widespread and abundant sub-species of green peafowl, with multiple fragmented populations distributed throughout China, Thailand, Cambodia, and Vietnam. The phenotypic differences among sub-species mainly occur in facial skin patterning, crest feathers (length and patterning), plumage coloration, shade, and size of the birds (Jackson 2006). Field biologists have argued for the occurrence of additional sub-species based on outer morphology (Lin 2010). However, the genetic distinctiveness of these phenotypes and geographically isolated populations has never been comprehensively assessed (Gu and Wang 2021).

Contemporary conservation efforts rely upon the maintenance of Studbooks that document the population’s geographic location for breeding purposes (European Conservation Breeding Group 2023). Nevertheless, such approaches have limited success in incorporating confiscated individuals of unknown genetic identity and could favor the admixture of distinct gene pools, thus lessening the genetic distinctiveness of resident populations. Hence, non-genetics-based approaches have limited or no utility in the conservation of resident populations. Spalding or emerald peafowl resulting from crosses between green and blue peafowl (*Pavo cristatus*, Linnaeus 1758) is produced by private breeders for aesthetic purposes and may appear phenotypically identical to the green peafowl (Du et al. 2020). Such hybrids, most often of unknown ancestry, cannot be flagged as such without genetic tools and can potentially affect breeding stocks and lead to hybridization between species either in captivity or when the hybrids are unintentionally released in the wild. Hence, a comprehensive geographic sampling throughout the native range of the wild green peafowl and recording of detailed sub-species discriminating phenotypic characters (such as crest feathers, plumage coloration, and shade) along with genetic/genomic sequencing of the same individuals is a fundamental prerequisite for establishing conservation programs.

Conservation policies for species with a fragmented distribution across multiple countries require an international outlook that can balance national economic and developmental priorities. In the case of Southeast Asia, conservation policy is conceived either at the international level (Convention on Biological Diversity), regional level (Association of Southeast Asian Nations), or national level (see **Table 1** for more details). Implementing these policies and monitoring outcomes depends on each country’s priorities. For instance, China’s Red list of vertebrates classifies the green peafowl as "critically endangered (CR)" while the IUCN as only "endangered (EN)". However, due to limited information on population genetic structure, the green peafowl has not been assessed at the sub-species level. Although the total number of mature green peafowl individuals is estimated at around 20,000, a large proportion may be from a single sub-species, likely *P. m. imperator*. Hence, the other two sub-species (*P. m. spicifer* and *P. m. muticus*) may have very few individuals and would require urgent attention for their protection. Moreover, *P. m. muticus* conservation is important for reintroduction in their native ranges of peninsular Malaysia.

**Table 1:**
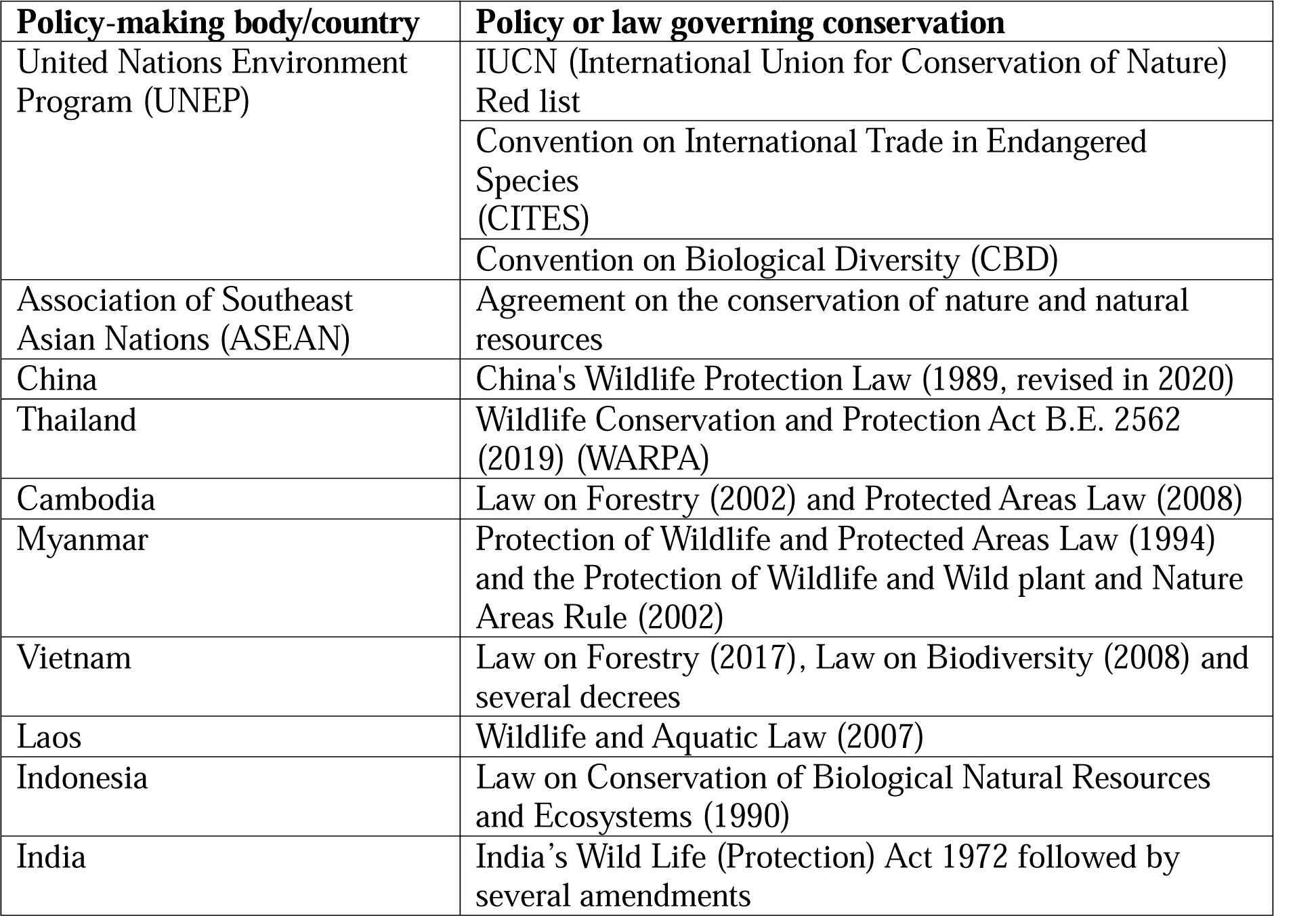
Conservation policies governing Southeast Asian biodiversity and wildlife protection.

Conservation initiatives for the green peafowl have focused on identifying its interactions with humans and their impact on the endangerment of this species (Dong et al. 2021). Habitat loss and fragmentation have wiped populations from the Malay Peninsula, yet individuals from this area still occur in protected parks, zoos, or museums, thus representing valuable resources for future reintroduction (Conway 2003, 2011; Fraser and Wharton 2007). To ensure that the distinctiveness of wild populations is retained in captive breeding programs, their genetic diversity and structure must be considered in population management (Wharton 2008). Historical demographic processes are often a major driver of current levels of genetic diversity in animal species and can provide insights into their response to future climatic fluctuations (Kozma et al. 2016; Song et al. 2020). The subsequent genetic screening of captive individuals might help identify their source populations in compliance with current tenets of ex-situ conservation (Wharton 2008). Hence, genetic and genomic tools can be important in green peafowl conservation management (Höglund et al. 2019; Segelbacher et al. 2021). We envisage that genetics can play a meaningful role in captive breeding programs concerning the wild reintroduction of green peafowl.

Based on these considerations, our objectives are:

1. To evaluate the occurrence of genetically distinct populations of green peafowl using genomic datasets.
2. To assess whether distinct populations experienced different historical bioclimatic events shaping their demographic histories.
3. To estimate the population split times.
4. To quantify the magnitude of genetic variation along the native ranges to help define management units (MU) and provide ancestry-defining markers for these populations.

Both morphologic and genetic evidence will be key to defining Evolutionarily Significant Units (ESU) (Conner and Hartl 2004) or Management Units (MU) (Palsbøll et al. 2007) for effective conservation management of the green peafowl.

## Materials and Methods

### Data curation

Recently two *P. muticus* de novo genome assemblies were published, and 52 whole genomes were re-sequenced (Dong et al. 2021; Zhang et al. 2022). We downloaded paired-end reads and obtained the associated metadata for one *P. cristatus* (from NCBI) and 52 *P. muticus* individuals (38 individuals from NCBI and 14 from CNGBdb): **Supplementary Table 1**.

### Genomic data analysis and variant calling

We selected the most recent chromosome-level genome (Zhang et al. 2022), which is comparatively more contiguous than the other based on its assembly metrics. We indexed the reference genome and mapped all the downloaded paired-end reads to the unmasked genome using the BWA MEM mapper (Li 2013) with default settings. The details of the percentage of mapped reads and coverage are provided in **Supplementary Table 2**. Genomewide variants were called using the bcftools pipeline (Li and Barrett 2011). We applied quality thresholds like - C 50 (mapping quality normalization) -q 30 (mapping quality cutoff) -Q 30 (base quality cutoff) for mpileup and used multiallelic caller to obtain the all site vcf. We further filtered the vcf file by removing indels, applying the maf filter of 0.1, and reducing the missing data threshold to 90 percent.

### Population structure analysis

We used ANGSD version: 0.935-53-gf475f10 (Korneliussen et al. 2014) to call genotypes with stringent filtering criteria (-SNP_pval 2e-6 -minMapQ 30 -minQ 20 -minMaf 0.05 -uniqueOnly 1 -remove_bads 1 -only_proper_pairs 1 -trim 0 -C 50 -baq 1 -setMinDepthInd 10) to maintain the high quality of the data. The called genotypes were then used as input for PCAngsd (Meisner and Albrechtsen 2018) to perform Principal Component Analysis (PCA) and NgsAdmix (Skotte et al. 2013) for estimating individual admixture proportions. To determine the best K (expected number of genetic clusters or ancestral populations), 10 replicates per each K were run in NgsAdmix prior to calculating the Coefficient of Variation (CV) and deltaK (Evanno et al. 2005). The best K was K=3 using both methods (see **Supplementary Figure 1A-B**). The genomewide variant calls were used to assess structure using the fastSTRUCTURE program with 10 replicates each to find the best K (Raj et al. 2014). Similar to the results from NgsAdmix, the best K for fastSTRUCTURE results were found to be K=3 using Evanno method (Evanno et al. 2005) implemented in CLUMPAK (Kopelman et al. 2015). The PCA, NgsAdmix, and fastSTRUCTURE analysis were done separately (1) using only the 52 *P. muticus* individuals and (2) using the 52 *P. muticus* individuals along with the *P. cristatus* individual. In both cases, the best K was found to be 3.

To generate an SNP-based phylogenetic tree, we used the vcf file for 52 *P. muticus* and one individual of *P. cristatus* as an outgroup. The vcf file was filtered to remove missing data and it was pruned to remove sites in linkage. We first converted the vcf file to HapMap format, which was then used as input for the SNPhylo pipeline (Lee et al. 2014). The obtained alignment from the SNPhylo was used to find an appropriate model of evolution using modeltest-ng followed by phylogenetic tree inference using raxml-ng with 1000 bootstraps (Kozlov et al. 2019; Darriba et al. 2020). The obtained consensus tree was visualized using FigTree.

### Genetic diversity, divergence, and differentiation

We used BAM alignments of all the individuals to the reference genome of *P. muticus* to estimate Site Frequency Spectrum (SFS) by estimating genotype likelihoods using ANGSD. We applied multiple quality thresholds to restrict analyses to high-quality variants. We only used uniquely mapped and properly mate-mapped reads, excluding the bad (flag >=256) ones. Further, we restricted our variant sampling to the sites where mapping quality adjustment (-C 50), adjusting q-scores around indels (-baq 1), minimum mapping quality of 20 (-minMapQ 20), minimum base quality of 20 (-minQ 20), at least 75 percent of individuals considered (-minInd Nx0.75), depth of at least 10 bases at each site in each individual (-setMinDepthInd 10). The SFS was obtained using the realSFS module of ANGSD from the obtained SAF (Sample Allele Frequency) files.

The genetic diversity of each population was estimated using ANGSD -doThetas flag with the SFS obtained in the previous step as a reference using saf2theta module. The theta estimates were obtained using the thetaStat module for 50 KB non-overlapping windows. To obtain global diversity estimates, we calculated the mean of Watterson’s estimator (θ_w_) and Pairwise nucleotide diversity (θ_π_) of the 50 KB windows normalized by the number of sites in each window for each population, separately for autosomes, Z and W chromosomes.

We inferred pairwise 2D-SFS for the population pairs (Vietnam vs. Yunnan, Vietnam vs. Thailand, and Thailand vs. Yunnan) to obtain differentiation estimates. These 2D-SFS were indexed and used to calculate estimates of F_ST_ in per-base and for 50KB non-overlapping windows for each population pair. To get the global F_ST_ values, we calculated the mean of F_ST_ estimates for 50 KB windows of autosomes for each population pair. We used the pixy pipeline to calculate divergence estimates, i.e., d_XY_ (Korunes and Samuk 2021). The chromosome-wise vcf file was used for calculating θ_π_, d_XY,_ and F_ST_ using pixy for 50 KB non-overlapping windows.

To calculate haplotype statistics, we first phased all filtered chromosomal vcf files using Beagle 5.4 (Browning et al. 2021). We further subset the vcf for each population and chromosome. We used the R package “rehh” to filter variants to a MAF cutoff of 0.05 and calculate iHS statistics for each population (Gautier et al. 2017). To calculate xpEHH for each population pair, we compared the iHS values of the corresponding population pair. The obtained xpEHH estimates were then exported into text files for visualization in R. To identify outliers, we decided on the cutoff of 95 percentage quantile, i.e., the windows with the top 5 % highest values of the autosomal estimates of F_ST_ and d_XY,_ respectively. The obtained windows of F_ST_ and d_XY_ were then intersected to find regions of high differentiation and divergence.

### Demographic history reconstruction

The BAM alignments for each sample were used to generate a consensus sequence using samtools (Li et al. 2009) and bcftools (Li and Barrett 2011) with quality filters of -C50 -Q20 - q20. The consensus calls were converted to fastq format using vcfutils.pl vcf2fq with a quality filter (-Q) of 25 and a coverage filter greater than twice and less than one-third of the mean coverage. Resultant fastq files were converted to the psmcfa file using the fq2psmcfa command of the psmc (Pairwise Sequentially Markovian Coalescent) suite. We then used the psmcfa file to run psmc (Li and Durbin 2011) with options -N 30 -r 5 -t 5 -p 4+30*2+4+6+10. Pairwise pseudo-diploids constructed between individuals of different populations identify the historical splits by capturing the cross-coalescence between them. Hence, the time point at which the PSMC trajectory is vertical denotes the break in gene flow between these populations (Cahill et al. 2016; Song et al. 2017; Kishida 2017). For pseudo-diploid analysis, the fastq consensus files were converted to fasta using the seqtk fq2fa module. These fasta sequences for a pair of samples were merged using the seqtk mergefa module and then converted back to fastq format to generate a .psmcfa file using the fq2psmcfa module. Two different time intervals, i.e., -p parameters (-p "20+4*5" and -p "4+25*2+4+6") were chosen to get sufficient recombinations and resolution at a particular time point. The psmc files were evaluated for an adequate number of recombination events. The estimated trajectory of effective population size (N_e_) was plotted with a mutation rate (u) of 1.33e-09 per site per year (Wright et al. 2015) with a four-year generation time using psmc_plot.pl script of psmc.

### AIMs identification

To identify Ancestry Informative Markers (AIMs), we used vcftools.jl, part of openmendel julia utilities (Zhou et al. 2020; Ko et al. 2023). The autosomal phased vcf file of individuals from native populations (i.e., Vietnam, Yunnan, and Thailand) was used to rank the SNPs based on their ability to distinguish the ancestry. The SNP rank was used to select the best-ranked top 100 SNP. These 100 selected SNP’s were then used to run ten replicates of fastSTRUCTURE to find the best K (Raj et al. 2014). The best K using Evanno’s Method was K=3 (Evanno et al. 2005). The obtained clusters were observed for the assignment of the individuals in each cluster (**Supplementary Figure 2A-I**).

To understand the effect of demographic history on the AIMs panel, we assessed the temporal distribution of these top-ranked SNP’s, i.e., the timeline/TMRCA (Time to Most Recent Common Ancestor) bin to which these SNP’s contribute. PSMC was run for a representative individual from each of the three native populations with -d parameter, which gives the regions contributing to each time interval. We selected the top 10,000 ranked SNP and intersected them with the regions contributing to each time interval in PSMC. The number of these top-ranked SNP in each time interval was calculated for each representative population and visualized comparatively using a barplot to identify differences in the distribution.

## Results

### Green peafowl population structure and diversity

Genomewide patterns of genetic variability along the first principal component (5.83%) in the green peafowl captured the distinctiveness of individuals from Yunnan (southwestern China), Thailand, Cambodia, and Vietnam (see **Figure 1**). The second axis (3.49%) revealed the substantial variability within the samples from Yunnan and separated those samples from QHD (Qinhuangdao Wildlife Park, China). The samples from the Xinxing breeding base either clustered with Vietnam or formed a separate cluster in the PCA. Admixture analysis revealed that some samples (XNJ7, 8, and 11 when K=4) from the Xinxing breeding base (China), closely clustering in the PCA with those from Vietnam and Cambodia, were indeed admixed with them (see **Figure 2**). However, other samples from this captive population formed a separate cluster when K=4. Overall, QHD samples showed the highest level of genomic distinctiveness compared to the other populations. The YN1 individual also appears to have both QHD and Yunnan ancestry. PCA and admixture analysis, including a blue peafowl genome, suggest QHD samples are admixed (see **Supplementary Figure 3-4**). The SNP-based phylogenetic tree supports the population clusters identified in PCA and admixture (see **Supplementary Figure 5**). Estimates of autosomal genetic diversity are much higher in the QHD samples (θ_π_ of 0.00288) compared to the native *P. muticus* populations (θ_π_ of 0.00175 (Yunnan), 0.00183 (Thailand) and 0.00177 (Vietnam)) (see **Supplementary Table 3-4**). The native populations (θ_w_ of 0.00158 (Yunnan), 0.00170 (Thailand), and 0.00160 (Vietnam)), have a higher genetic diversity than the samples from the Xinxing breeding base (θ_π_ of 0.00163 and θ_w_ of 0.00145). Therefore, the genetic diversity across the native populations was comparable, although samples from Yunnan are known to have a slightly lower diversity (Dong et al. 2021).

**Figure 1:**
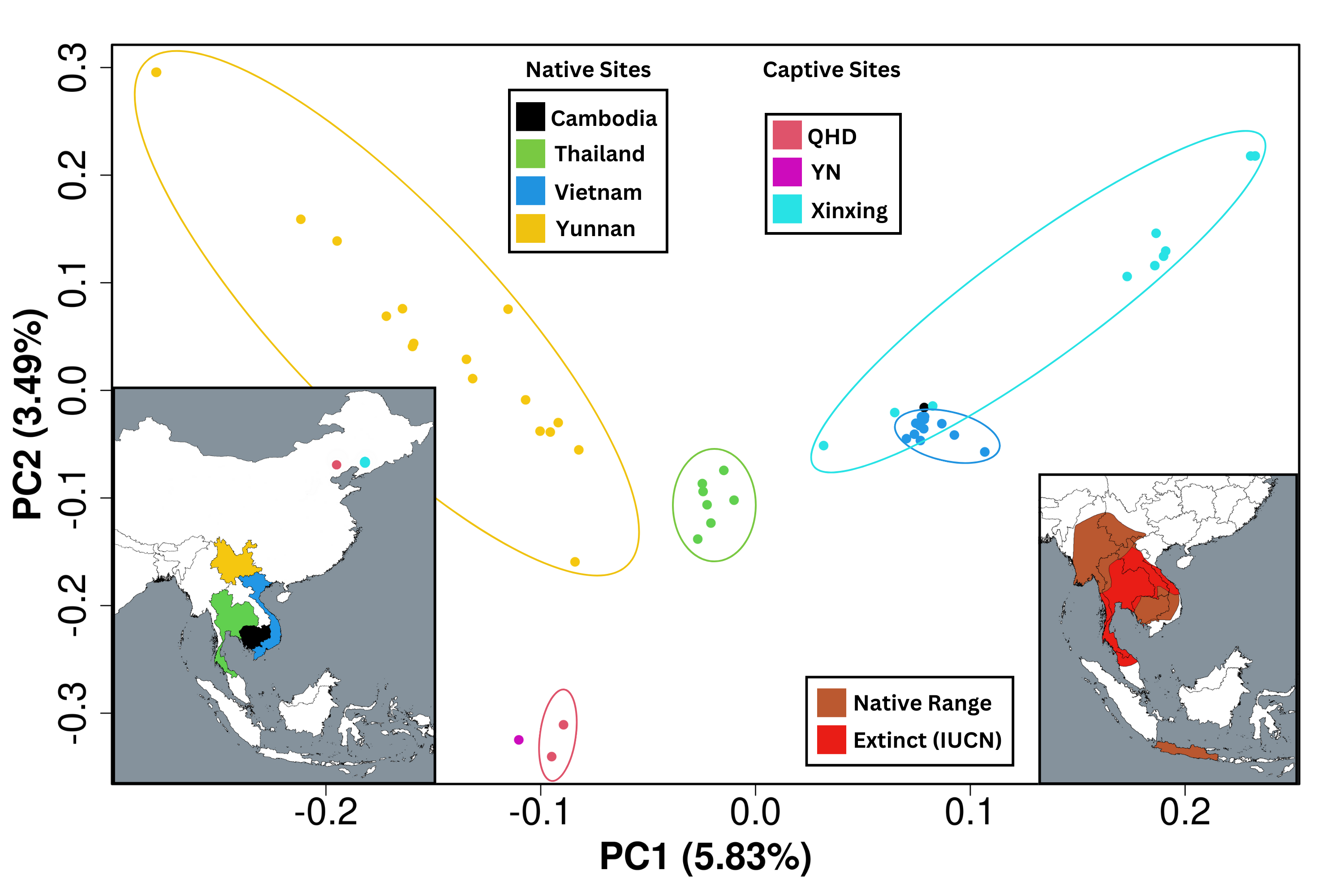
Principal component analysis (PCA) based on 1.88 million SNP’s illustrating the genetic variation in 52 individuals of *P. muticus*. Each dot represents an individual, whereas colors represent countries and captive populations. The map at the bottom left corner shows the countries where the species is presently found, while the bottom right corner shows the current and former range based on the IUCN database. PC1 explained 5.83% of genetic variation and separated samples from Yunnan (Yellow), Thailand (Green), Vietnam (Blue) plus Cambodia (Black), and Xinxing breeding base (Cyan) populations, with three individuals from Xinxing clustered among individuals from Vietnam and Cambodia. PC2 explained 3.49% of genetic variation and separated mainly QHD (Pink-red) and YN (Magenta) individuals from other populations.

**Figure 2:**
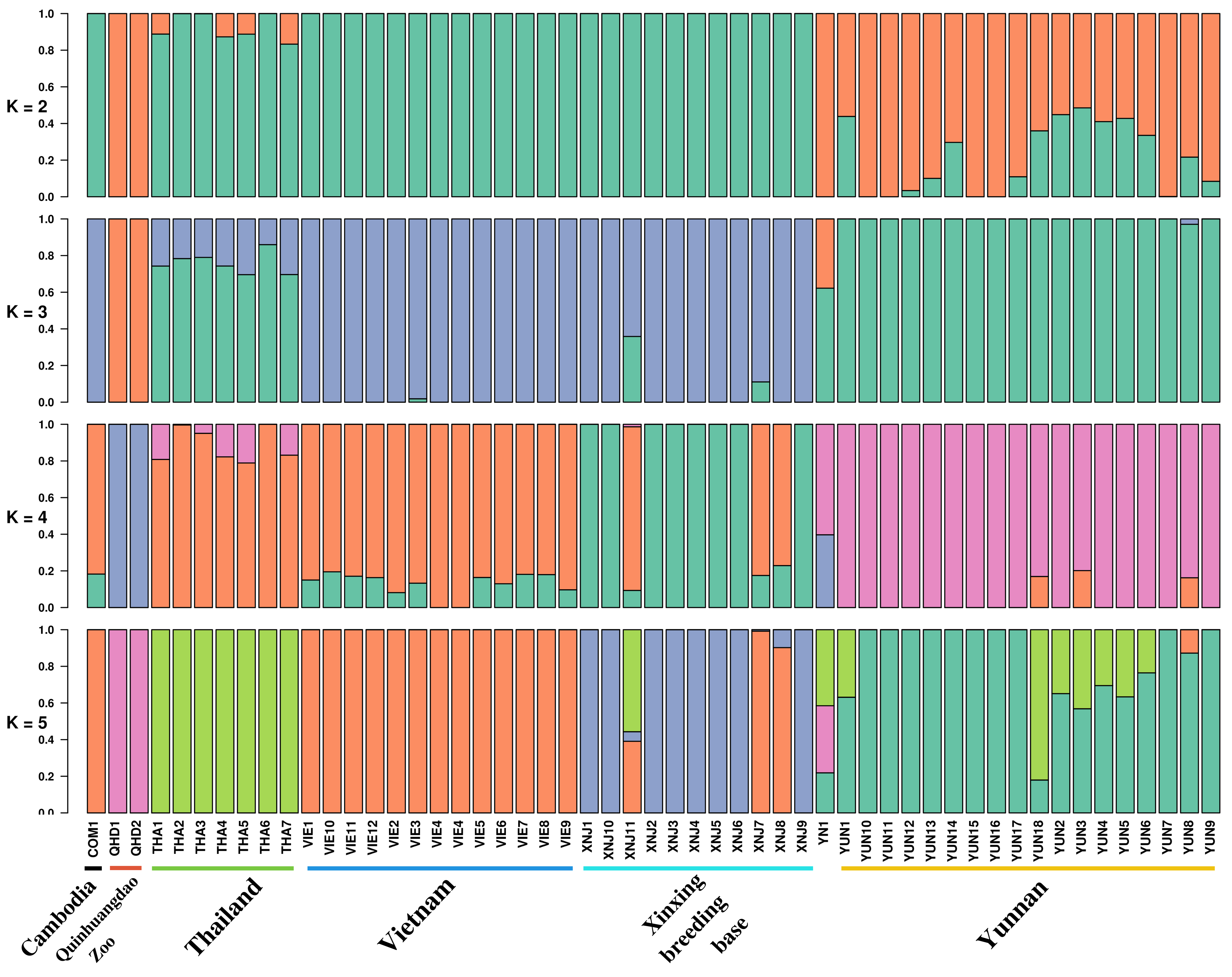
Admixture analyses and population clustering of *P. muticus* populations. Population structure inferred in NGSAdmix based on genotype information in 52 *P. muticus* individuals. The Coefficient of Variation (CV) and deltaK by Evanno’s method across log-likelihood values for ten runs at each K value (1 to 8) showed the best K=3. QHD individuals and YN1 form a unique cluster that shares ancestry with *P. cristatus* and YUN (Yunnan) samples (see **Supplementary Figure 4**). YUN and THA (Thailand) populations show close ancestry at K=3. In contrast, they separate at a further resolution of K. Individuals from VIE (Vietnam), COM (Cambodia), and XNJ (Xinxing breeding base) seem to form another cluster and show ancestry, which further separates at higher K values.

### Genetic differentiation

We considered all three population comparisons among native populations, i.e., Yunnan vs. Thailand, Vietnam vs. Yunnan, and Thailand vs. Vietnam, for calculating differentiation estimates (see **Supplementary Table 5**). The overall autosomal mean F_ST_ for Yunnan vs. Vietnam pair was higher (0.0995) than other pairs (i.e., Thailand vs. Vietnam = 0.0772 and Yunnan vs. Thailand = 0.0761). Even though levels of differentiation were higher in Yunnan (n=18) vs. Vietnam (n=12) pair, it has fewer fixed sites (3) than the other two pairs (Thailand vs. Vietnam = 12 and Yunnan vs. Thailand = 14), probably due to the smaller sample size of Thailand (n=7). Although the number of fixed sites is known to be vulnerable to sequencing errors or the number of samples used, the genomewide levels of differentiation mirror the geographic distribution. Additionally, the relatively higher F_ST_ between Yunnan and Vietnam supports the two distinct clusters identified by admixture analysis. The window-wise outlier scans for highly differentiated regions (i.e., windows with top 5% F_ST_ and D_XY_) across population pairs identified candidate loci that might be under selection (see **Supplementary Table 6**). Using the additional criteria that haplotype statistics (i.e., xpEHH) should also show a clear signal, we considered these regions as putatively under selection. The candidate region identified in the Yunnan-Vietnam comparison is on chromosome 4 (**Supplementary Figure 6A**). Regions on chromosomes 10, 20, and 25 are candidates in the Yunnan-Thailand comparison (**Supplementary Figure 6B-D**). The Thailand-Vietnam comparison has candidate regions on chromosomes 4 and 25 (**Supplementary Figure 6E-F**). The chromosome-wise results of outlier scans are provided in **Supplementary File S1-3.**

### Demographic histories

The historical demographic trajectories of all the green peafowl individuals have a comparable trend of population size expansion and contraction coinciding with the inter-glacial events (see **Figure 3A**). However, unlike all others, the two QHD individuals have a drastically different demographic trajectory since their population experienced expansion during the mid-Pleistocene (**Supplementary Figure 7**). This pattern of population expansion resembles the blue peafowl population expansion (see **Supplementary Figure 8**). Moreover, the trends reconstructed based on these QHD samples differed from each other since 1000 KYA (1 MYA) in both magnitude and trajectory. The other closely related inland populations (Yunnan, Thailand, Vietnam, and Xinxing breeding base) have very similar trajectories until MIS9 (Marine Isotope Stage 9) inter-glacial period (**Figure 3B** and **Supplementary Figures 9-12**). Nonetheless, the demographic trajectories of these populations began to diverge after the MIS9 inter-glacial period. The similarity of the demographic trajectories mirrored the genetic structure and biogeographic pattern (see **Figure 1-3** and **Supplementary Figures 7-12**).

**Figure 3:**
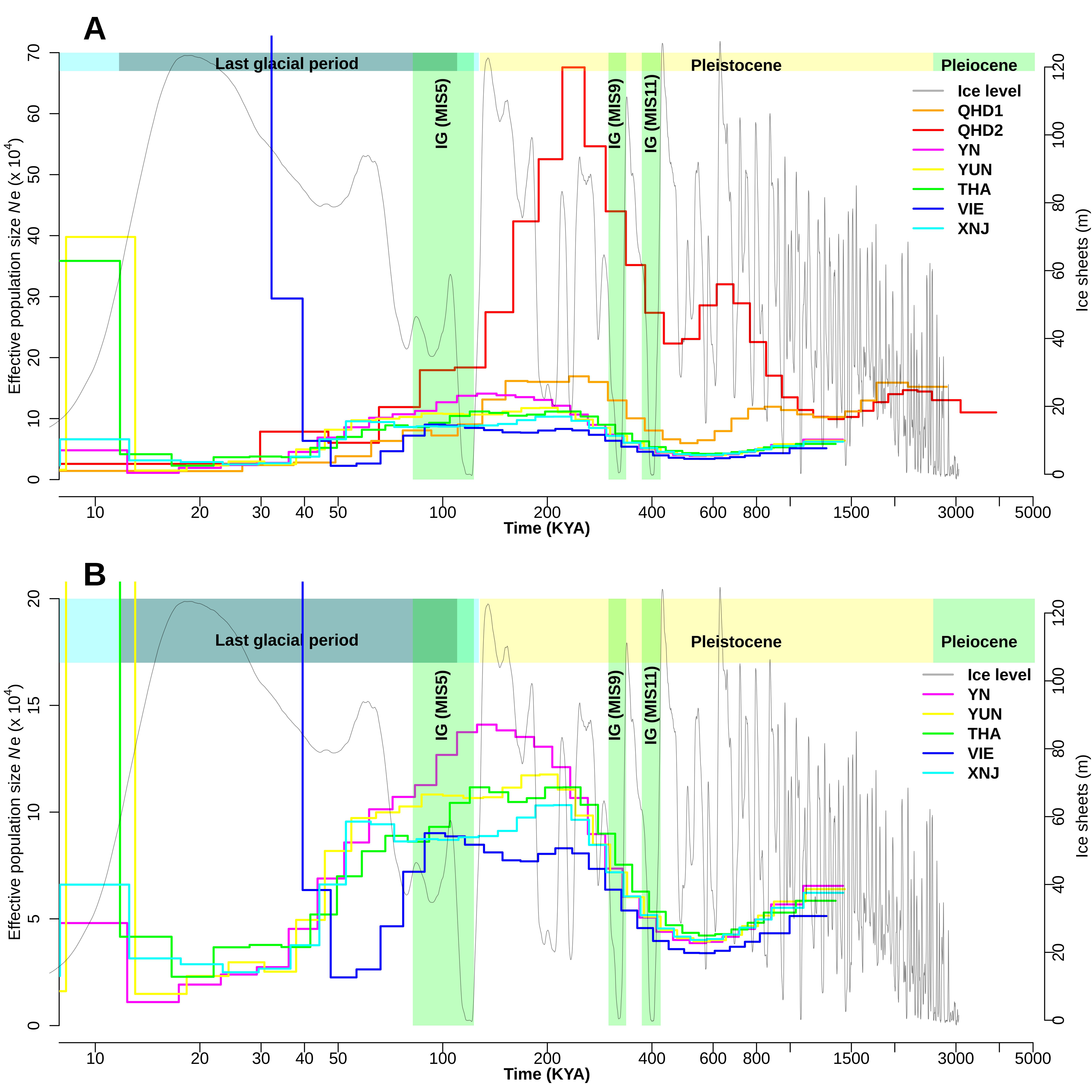
Historical demographic trajectories of *P. muticus* populations. A) Trajectories depicting demographic histories of *P. muticus* populations. QHD individuals show completely different trajectories compared to all others. B) Trajectories depicting demographic histories of closely related *P. muticus* populations.

### Population split times

The split times estimated by pseudo-diploid of all native green peafowl individuals compared with blue peafowl ranged between 3000 KYA to 1000 KYA (3-1 MYA), whereas QHD individuals with blue peafowl ranged between 3000 KYA to 600 KYA (3-0.6 MYA). These results suggest all native green peafowl individuals split from blue peafowl much earlier than the admixed QHD individuals (See **Supplementary Figure 13**). The split times estimated between QHD individuals and native green peafowl individuals ranged between 2000 KYA to 500 KYA (2-0.5 MYA) and suggest an earlier split from native green peafowl individuals, possibly due to blue peafowl admixture. (See **Figure 4**). The split times of the admixed YN occurred earlier than those of the inland populations, which are posterior to MIS9. The split between the native green peafowl individuals (Yunnan, Thailand, and Vietnam) coincided with the MIS9 inter-glacial period, i.e., ∼ 300 KYA, and are latest to split among the other groups. These split time scenarios suggest the vicariance between green peafowl individuals during the interglacial period during MIS9, possibly due to rising sea levels leading to loss of connectivity between populations (See **Figure 4**). The SNP-based phylogenetic tree also supports this step-wise pattern of population splits (see **Supplementary Figure 5**).

**Figure 4:**
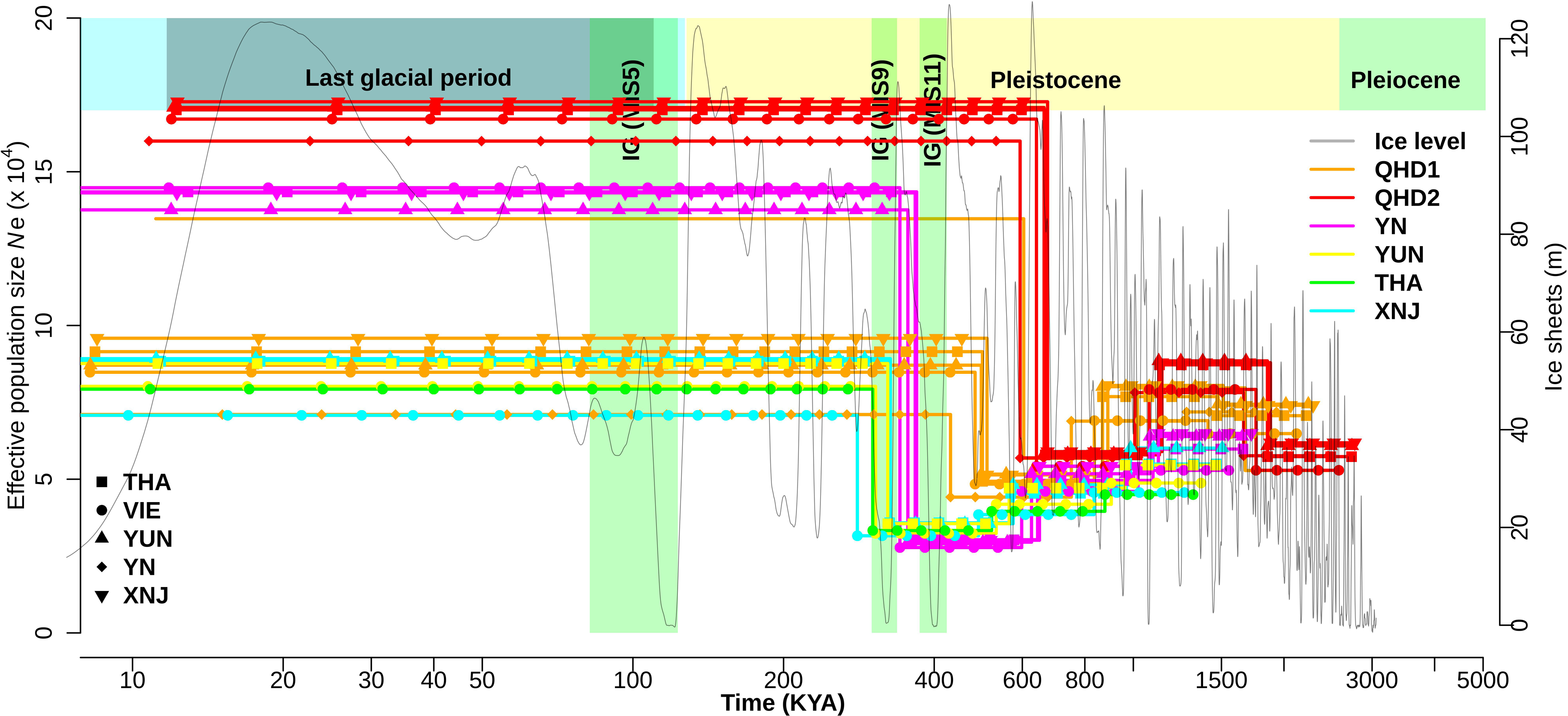
Split time estimates based on pseudo-diploids between pairs of individuals of different *P. muticus* individuals. Colors represent one individual from respective populations, whereas line types represent the other individual in a pair. QHD2 is the first to diverge from other populations, followed by QHD1, with other closely related populations showing a similar timeline for the split.

### Ancestry informative markers (AIMs)

The top-ranked 100 ancestry-defining SNPs can assign the green peafowl individuals to their respective geographic populations. The fastSTRUCTURE results obtained using these top 100 SNPs placed the individuals (native and captive) in their respective population clusters, confirming their application in population assignment (**Supplementary Figure 14**). Further, the distribution of the top 10,000 ranked ancestry-defining SNPs contributed most to the TMRCA bins of PSMC belonging to the 45-50th time intervals which correspond to the period around 300 KYA to 1000 KYA (0.3 to 1 MYA) (**Supplementary Figure 15**). The distribution of top-ranked SNPs in these older time bins suggests these markers capture ancestral haplotype structure. However, in the individual from Thailand, a large proportion of the ancestry-defining SNPs are found in the recent time bins suggesting they are affected by population-specific homozygosity tracts. Hence, the AIMs identified in our study capture the differences in the demographic history.

## Discussion

The green peafowl is an endangered species due to anthropogenic habitat degradation in the recent past (Dong et al. 2021). Continued threats to its habitat and conflicting socioeconomic priorities in Southeast Asia have made the green peafowl an iconic example of the perils faced by other species endemic to this region. The ongoing conservation efforts require habitat protection, which might, nonetheless, conflict with developmental goals. The definition of conservation units is crucial to implementing successful conservation programs. The genomic revolution has enabled high-throughput techniques to generate massive amounts of genomic data to assess population structure and reconstruct demographic histories in multiple species of conservation interest (Talla et al. 2023). However, conservation policies for species whose distribution spans multiple countries (i.e., transboundary species) are challenging to implement (Mason et al. 2020). The Southeast Asian countries encompassing the native range of the green peafowl lack legislative recognition for any form of ESU or MU. Formal recognition of such conservation units across Southeast Asia will be beneficial for promoting effective biodiversity conservation. The lack of studies investigating population structure in Southeast Asian species may be a major hurdle in defining ESU/MU. Hence, identifying population structure and its conservation relevance is an important step to spur a change in the policy.

Using previously published re-sequencing (Jaiswal et al. 2018; Dong et al. 2021; Zhang et al. 2022) datasets, we identified population structure corresponding to geographic distribution within the green peafowl. These populations have different demographic histories, allegedly resulting from different responses to interglacial events. Moreover, the coalescence-based split time analyses and SNP-based phylogenies suggest breaks in gene flow have established the genetic distinctiveness of these populations. Estimates of genetic differentiation between populations and ancestry-defining markers further support delineating three populations, i.e., Yunnan, Thailand, and Vietnam. Overall, our results evidence the occurrence of a marked population structure in the green peafowl, which is concordant with their geographic distributions.

More specifically, the individuals of Vietnam and Cambodia form a distinct cluster overlapping with the cluster of Xinxing breeding base in the PCA (**Figure 1**), suggesting a separation from other populations. In the admixture analysis at a K value of three, Vietnam, Cambodia, and Xinxing breeding base individuals show a single cluster, whereas Yunnan and QHD form separate clusters (**Figure 2**). However, at a K value of four, the admixture results mirror genetic variation explained in the first principal component, as three individuals of the Xinxing breeding base are similar to the Vietnam population while the remaining individuals are distinct. The SNP-based phylogeny also places these three individuals in the Vietnamese clade, while the other Xinxing breeding base individuals form a separate clade (**Supplementary Figure 5**). Even though the other individuals from the Xinxing breeding base are distinct from these three individuals, their closest relationship is with Vietnam in the phylogeny and admixture analysis. Hence, Vietnam appears to be the most likely source of all the Xinxing breeding base captive individuals.

The individuals from Thailand form a cluster between Vietnam and Yunnan populations in the PCA. At K=3 in the admixture analysis, Thailand appears to have shared ancestry with Yunnan and Vietnam while forming a separate cluster at K=5, consistent with their intermediate clustering in the PCA. The SNP-based phylogeny also corroborates this intermediate placement of Thailand individuals between the Vietnam and Yunnan clades. The green peafowl distribution in Vietnam is restricted to the South-central part adjacent to Cambodia (Sukumal et al. 2015). Therefore, Thailand’s population occurs geographically between the populations from Yunnan and Vietnam/Cambodia (Sukumal et al. 2020). The samples from Yunnan form a dispersed cluster in the PCA along both PC1 and PC2. The admixture analysis hints toward the substructure within the samples from Yunnan. Specifically, individuals YUN1-6 (museum samples) and YUN18 appear to have both Yunnan and Thailand ancestry. These seven samples are sampled from the Southern part of Yunnan and form a separate clade in the SNP-based phylogeny. Especially YUN18, sampled from the border of Yunnan (China) and Thailand, appears to have a greater Thai ancestry and forms a separate clade in the SNP-based phylogeny. Hence, the genetic structure matches the geographic distribution range of the green peafowl.

Three individuals from Qinhuangdao Wildlife Park (QHD1-2 and YN1) cluster separately from all other populations in PCA, suggesting the divergent origin of these individuals. In the PCA, blue peafowl clusters with QHD and YN1 individuals (see **Supplementary Figure 3**). Admixture analysis shows that QHD1 and QHD2 have differing levels of blue peafowl ancestry. However, the blue peafowl ancestry in YN1 is not evident in the admixture results (see **Supplementary Figure 4A-B**). In contrast, QHD ancestry in YN is picked up when blue peafowl is not included in the analysis (see **Figure 2**). The divergent clustering of these individuals with blue peafowl suggests inter-species hybridization with blue peafowl. However, another possibility is they may originate from Burman (*P. m. spicifer*) green peafowl which may have blue peafowl ancestry due to its geographic proximity to the range of the blue peafowl.

Changes in sea level due to glacial events lead to either increased or decreased connectivity between islands due to exposed or submerged land bridges. The Southeast Asian islands are vulnerable to these sea level changes and have experienced several recurrent events resulting in landmass reconfigurations (Voris 2000; Woodruff 2010; Salles et al. 2021). The availability of optimum habitat heavily depends upon changes in landscape morphology and is a driver of population dynamics. The increase in sea level during the MIS9 inter-glacial separated the green peafowl populations on different islands. These isolated populations either expanded or contracted based on resources available within their range. Our analysis identifies divergent demographic histories for the green peafowl populations (see **Figure 3**), beginning in the MIS9 inter-glacial. The pseudo-diploid analysis also identifies population splits between the green peafowl populations during the MIS9 inter-glacial (see **Figure 4**). Based on this co-occurrence of increased sea levels with changes in demographic history and population splits, we argue that these populations were separated around 300 KYA due to the inter-glacial.

The estimated split between blue and green peafowl ranges from 3000-1000 KYA (3-1 MYA), concordant with previous phylogenetic estimates (Chen et al. 2021). The split between blue and QHD peafowls is much later than the blue and green peafowl split but earlier than the inter-populations splits of green peafowl (**Supplementary Figure 13**). The demographic histories of QHD individuals are strikingly divergent from those of other populations and different from each other, pointing to the distinct origin of these two individuals (**Supplementary Figure 7**). Additionally, split-time analyses suggest an earlier split time for QHD2 compared to QHD1, further establishing the distinct origin of QHD samples. The greater blue peafowl ancestry in QHD2 might be responsible for the most distinct demographic trajectory and earliest split among the green peafowl. Comparatively lower blue peafowl ancestry in QHD1 and YN1 might explain the intermediate split times and demographic histories.

Management Units (MU) have been defined as demographically independent populations whose population growth depends mainly on the local population history (Palsbøll et al. 2007). Since the green peafowl populations have divergent demographic histories, they are good candidates for defining distinct MU. In addition to the distinct demographic histories since the last inter-glacial, the inter-population differentiation between native green peafowl populations (Yunnan, Thailand, and Vietnam) ranges from 0.07 to 0.1. The most differentiated population pair is Vietnam vs. Yunnan, which are geographically isolated due to localized range extinction (**Figure 1**). Using autosomal SNPs, we could identify ancestry-defining markers for the three native green peafowl populations. These AIMs correctly assign individuals to the respective populations and can reconstruct the population structure demonstrating their utility and serving as evidence for defining distinct units. Differentiation-based outlier scans identified a few candidate regions that were further prioritized using population genetics and haplotype-based statistics. These putatively selected loci might reflect local adaptation and must be verified using extensive sampling. The divergent demographic histories, inter-population genetic differentiation, and population structure provide concordant evidence to consider these populations as distinct MU.

Our results highlight the genetic uniqueness of green peafowl populations from Southeast Asia. Based on the geographic locations, the individuals from Yunnan, Thailand, and Vietnam-Cambodia are within the reported range of *P. m*. *imperator* and might represent the population substructure within this sub-species (Delacour et al. 1977; Sukumal et al. 2020). The previous report of a lack of population structure within green peafowl might reflect restricted sampling (Dong et al. 2021). A major limiting factor to assessing the genetic diversity, differentiation, and population stratification completely is the lack of range-wide genomic sampling of the green peafowl including regions like North-East India and Myanmar for the *P. m. spicifer* and Malay peninsular region, and Java for *P. m. muticus*. Hence, distribution-wide genetic and morphological profiling focusing on sub-species defining characters such as facial skin patterning, crest feathers (length and patterning), plumage coloration, shade, and size of the birds is a must for effectively conserving green peafowl. In addition to morphological profiling, other metadata, such as sex, developmental stage, geographic location, and the captive or native origin of the sample, should be recorded along with the genomic data. The availability of such metadata facilitates follow-up studies that can provide a comprehensive understanding of the species’ dynamics.

Furthermore, green peafowl being a transboundary species, cooperative data sharing between the nations throughout the range will play a constructive role, and associated nations should form an consortium/organization to expedite this effort. Intentional or unintentional hybridization between blue and green peafowl and release in the wild is another problem (Gu and Wang 2021). The presence of inter-specific hybrids in captivity or wild can be a threat to the surviving native populations. Due to the phenotypic resemblance of the hybrids to the green peafowl, misidentification and re-introduction into wild or captive populations can erode the genetic composition. To distinguish these inter-specific hybrids resembling green peafowl from sub-specific forms like *P. m. spicifer*, the blue peafowl also needs to be sampled from the potential overlapping ranges, i.e., North-East India, Bangladesh, and Myanmar, which will allow us to understand the natural hybridization scenario (if any) between the two species and effectively conserve the green peafowl and its natural populations.

Hence, we propose the following actions:

1. Range-wide population survey to estimate genetic diversity and levels of differentiation between populations, construct Ancestry Informative Marker (AIM) panels, evaluate levels of inbreeding and mutation load, identify adaptive genetic variation and its association with phenotypes.
2. Use genetic markers to assess the ancestry and possible admixture of captive individuals in wildlife parks and breeding centers as well as confiscated individuals. The captive breeding programs should pursue the maintenance of population genomic distinctiveness.
3. Museum samples can be compared with modern samples to assess historical structure, diversity, distribution of the native populations and can greatly aid in identifying captive individuals from locally extinct populations (Dong et al. 2021; Ernst et al. 2022). These data can be an important resource for planning reintroduction programs.

## Conclusion

Our study provides multiple evidences to establish the presence of atleast two or more genetically distinct geographically isolated populations that can be defined as separate management units (MU). We provide ancestry informative markers (AIMs) to genetically distinguish the populations from Yunnan (China), Thailand and Vietnam. Although these markers are specific for this geographic location, they could supplement future marker panels based on range wide sampling. For the future, we recommend linking the observed morphological diversity with genetic distinctiveness among the green peafowl populations. While we characterized population structure and differences in demographic history, systematic characterization of phenotypic differences between these groups would be an important next step for evaluating the conservation implications. Concerted changes and shared policies for identifying threatened habitats and managing captive populations will benefit from this information. Local extinction events that result in the loss of adaptive variation can be prevented or remediated through reintroduction programs.

## Availability of data

All datasets used in this study are compiled from public repositories. The scripts and associated data from the analysis are available here: https://github.com/Ajinkya-IISERB/Pavo/Conservation/.

## Acknowledgment

We thank the Ministry of Human Resource Development for awarding a fellowship to ABP. The Department of Biotechnology, Ministry of Science and Technology, India (Grant no. BT/11/IYBA/2018/03) and Science and Engineering Research Board (Grant no. ECR/2017/001430) provided computational resources (i.e., Har Gobind Khorana Computational Biology cluster).

## Author contributions

ABP analyzed the genomic data and generated all the results. ABP and NV wrote the manuscript.

## Supplementary figures

**Supplementary Figure 1: (A)** Coefficient of variation (CV) for different K values in NGSAdmix admixture analysis. **(B)** DeltaK by Evanno method implemented in CLUMPAK.

**Supplementary Figure 2: (A-I)**: fastSTRUCTURE results inferred using top 100 ranked Ancestry informative markers (AIMs).

**Supplementary Figure 3**: Principal component analysis (PCA) for one *P. cristatus* and 52 *P. muticus* individuals. Each dot represents an individual, and the colors represent the respective population or species.

**Supplementary Figure 4A**: Admixture analyses inferred in NGSAdmix for one *P. cristatus* and 52 *P. muticus* individuals for values of K=2 till K=5.

**Supplementary Figure 4B**: Admixture analyses inferred by fastSTRUCTURE for one *P. cristatus* and 52 *P. muticus* individuals for values of K=2 till K=5.

**Supplementary Figure 5**: Phylogenetic tree for 52 individuals of *P. muticus* with *P. cristatus* as an outgroup based on unlinked SNPs.

**Supplementary Figure 6**: Selection scans, i.e., F_ST_, d_XY_, θ_w,_ and θ_π_ and xpEHH panels for chromosomes with outlier regions. **(A)** Yunnan vs Vietnam **(B1-3)** Yunnan vs Thailand and (**C1- 2**) Thailand vs Vietnam.

**Supplementary Figure 7:** Historical demographic trajectories of Qinhuangdao Wildlife Park (QHD) individuals.

**Supplementary Figure 8**: Historical demographic trajectories for *P. cristatus* and representative individuals from *P. muticus* populations.

**Supplementary Figure 9:** Historical demographic trajectories of Yunnan, China (YUN) individuals.

**Supplementary Figure 10:** Historical demographic trajectories of Thailand (THA) and Cambodia (COM) individuals.

**Supplementary Figure 11:** Historical demographic trajectories of Vietnam (VIE) individuals.

**Supplementary Figure 12:** Historical demographic trajectories of Xinxing breeding base (XNJ) individuals.

**Supplementary Figure 13**: Split time estimates between *P. cristatus* and representative individuals from each population of *P. muticus* represented by solid lines. Colors and symbols represent the respective population of *P. muticus*.

**Supplementary Figure 14:** fastSTRUCTURE results for 53 peafowl inviduals inferred using top 100 ranked Ancestry informative markers (AIMs).

**Supplementary Figure 15**: Distribution of top 10000 ranked ancestry informative SNPs across PSMC free time intervals.

