## Supplementary figures and images for "Conservation implications of diverse demographic histories: the case study of green peafowl (*Pavo muticus*, Linnaeus 1766)"

### Supplementary_Figure1A.png

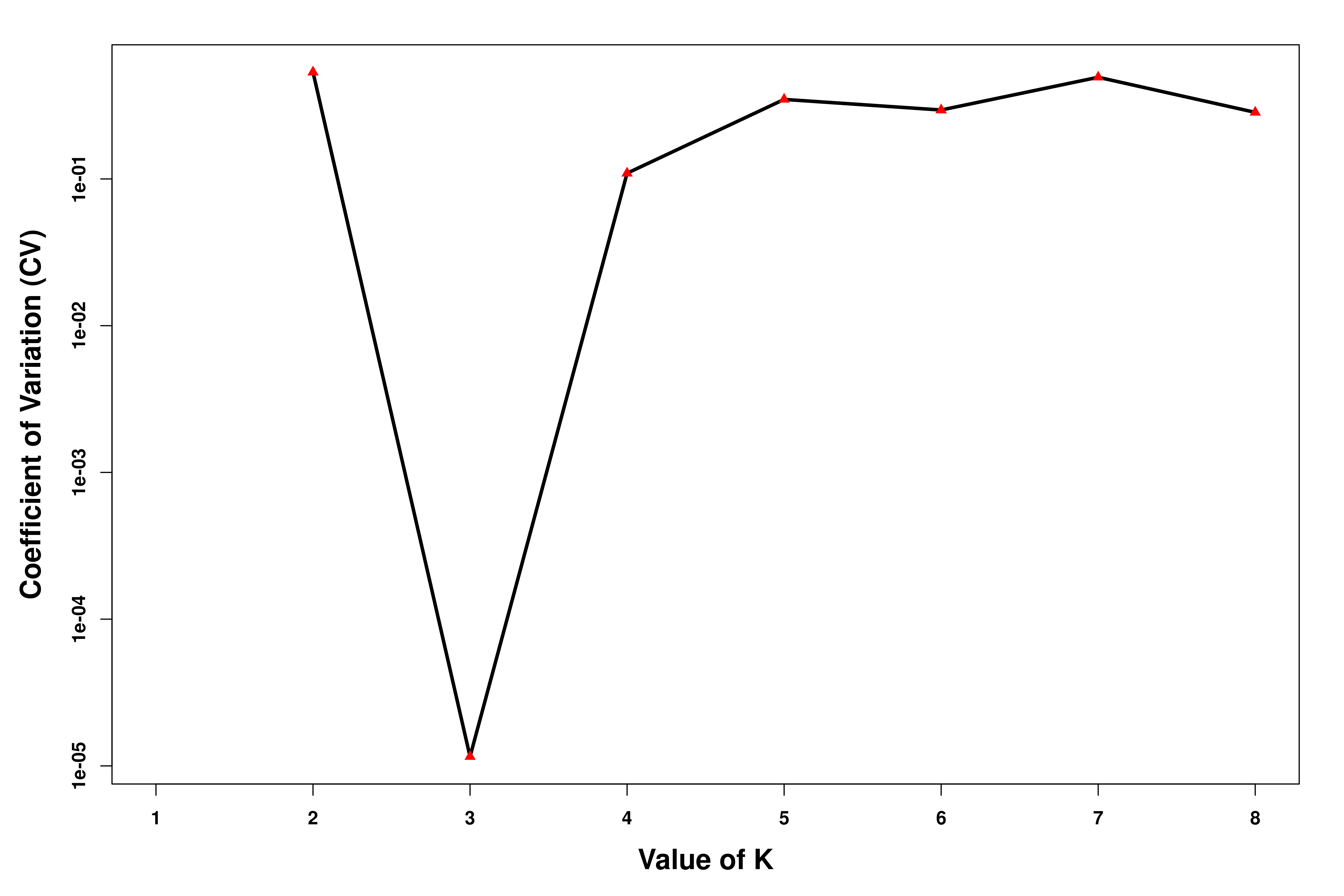

### Supplementary_Figure_1B.png

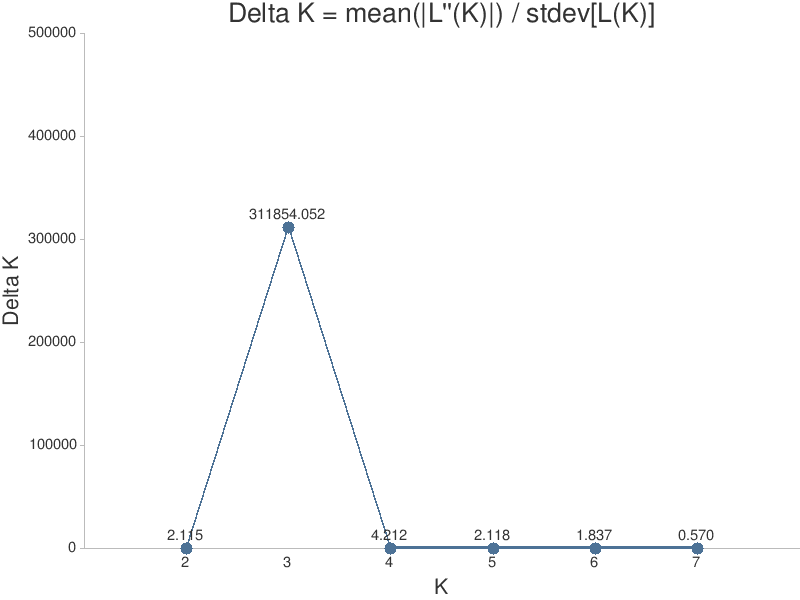

### Supplementary_Figure_2A.png

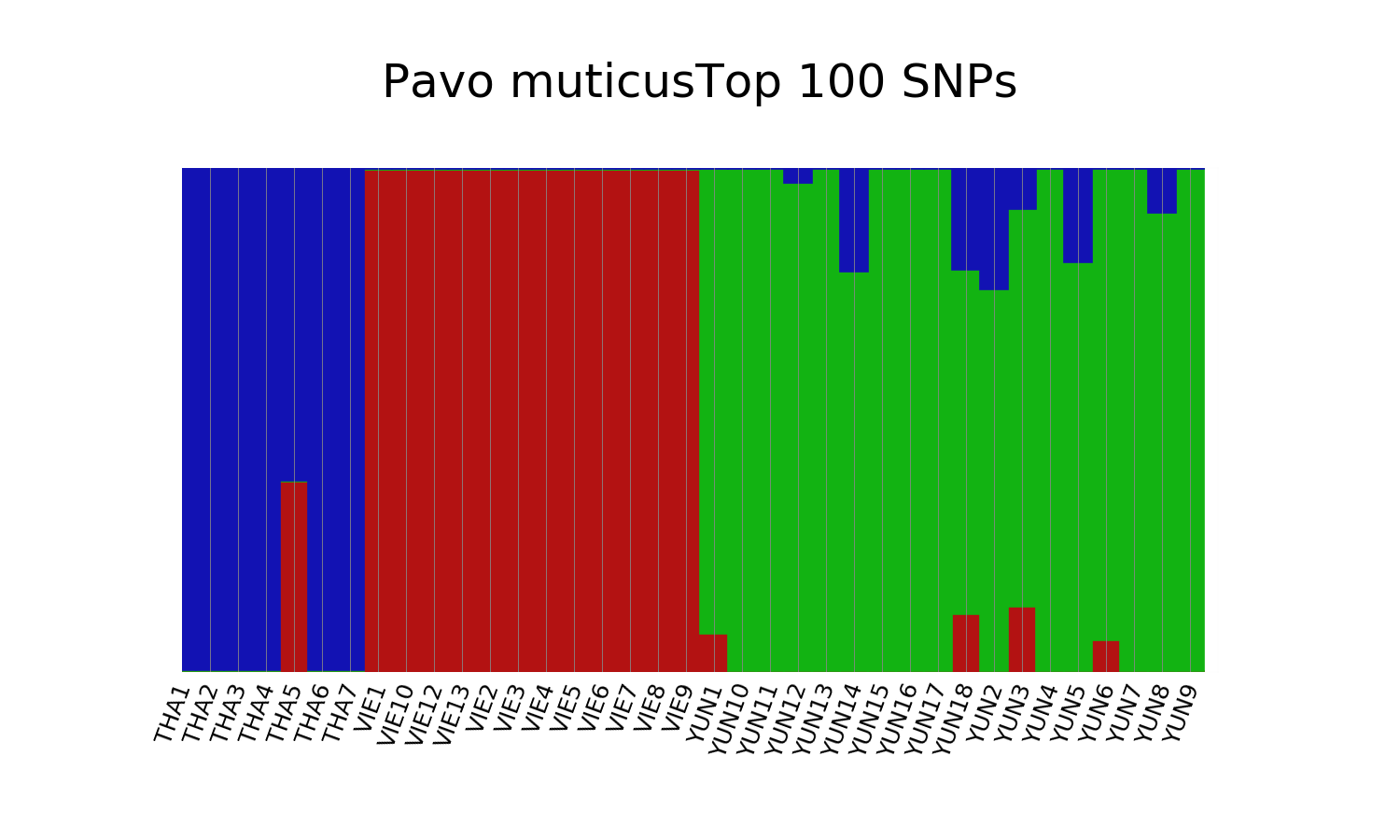

### Supplementary_Figure_2B.png

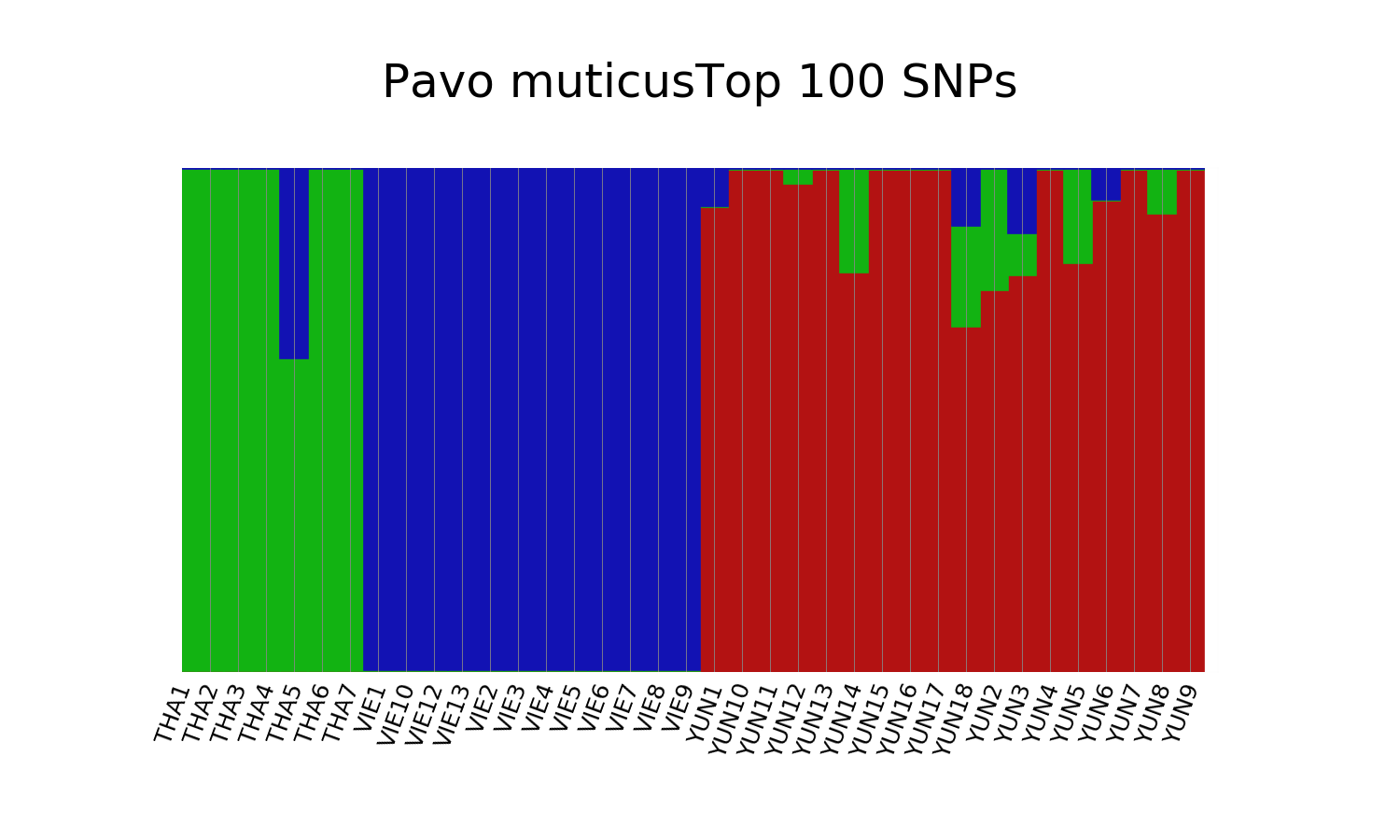

### Supplementary_Figure_2D.png

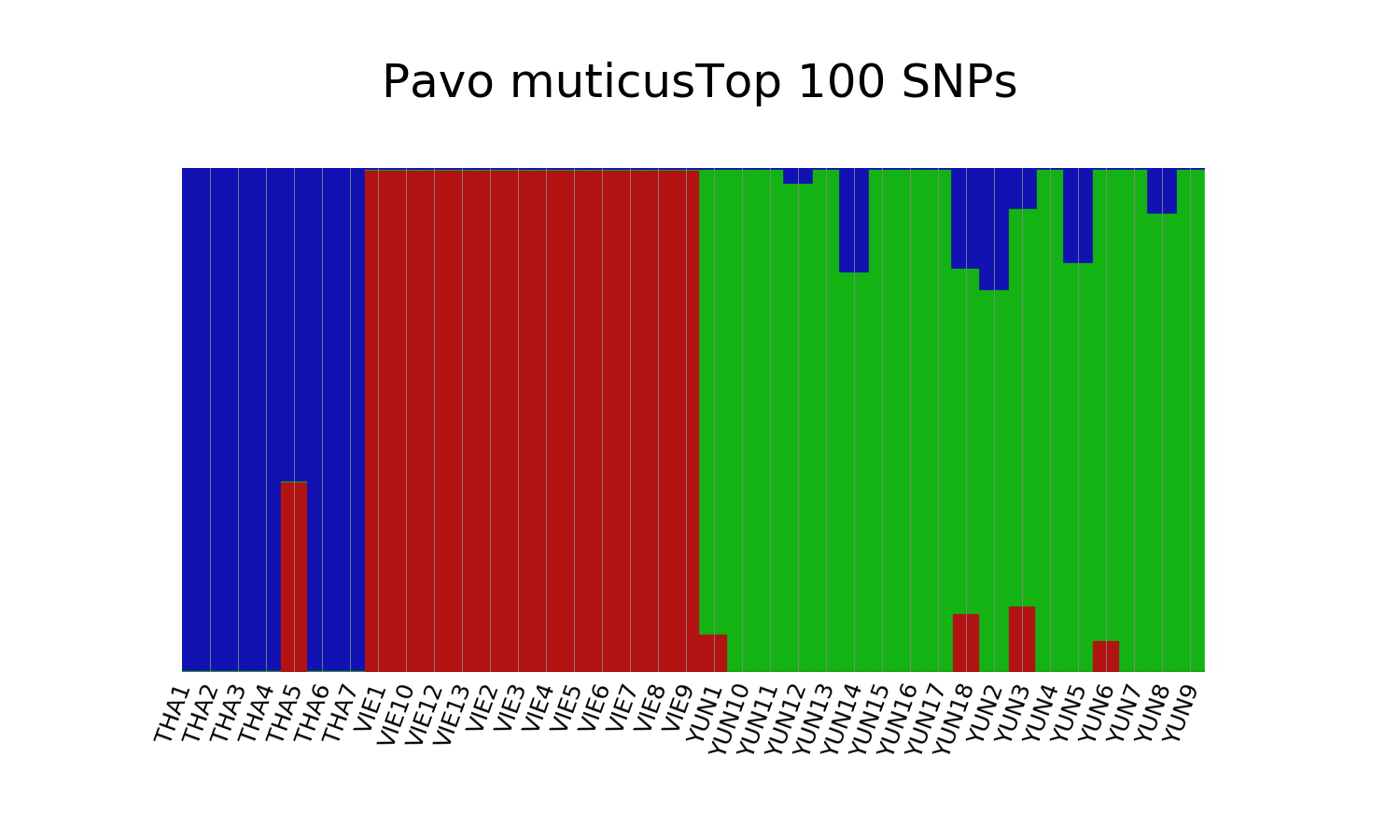

### Supplementary_Figure_2E.png

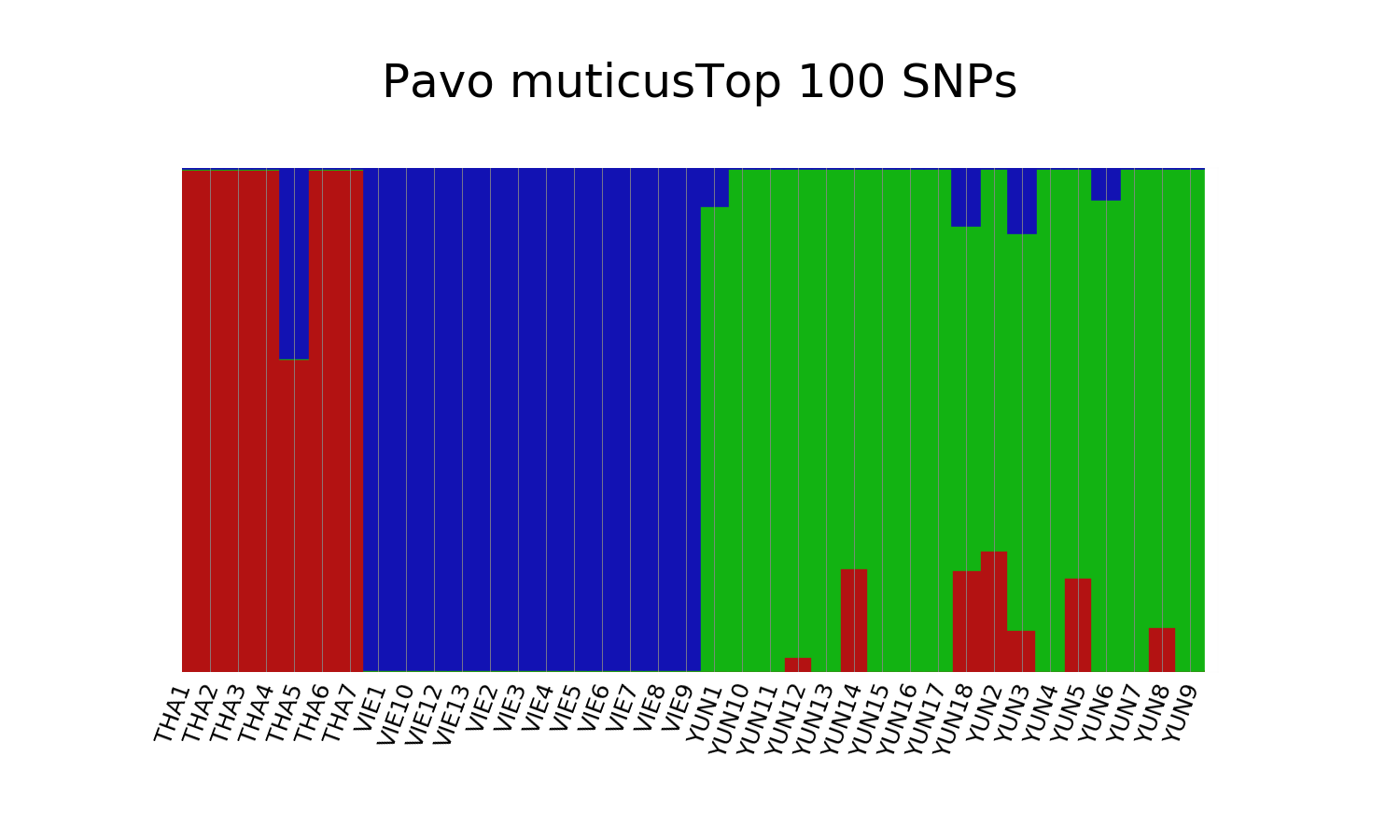

### Supplementary_Figure_2F.png

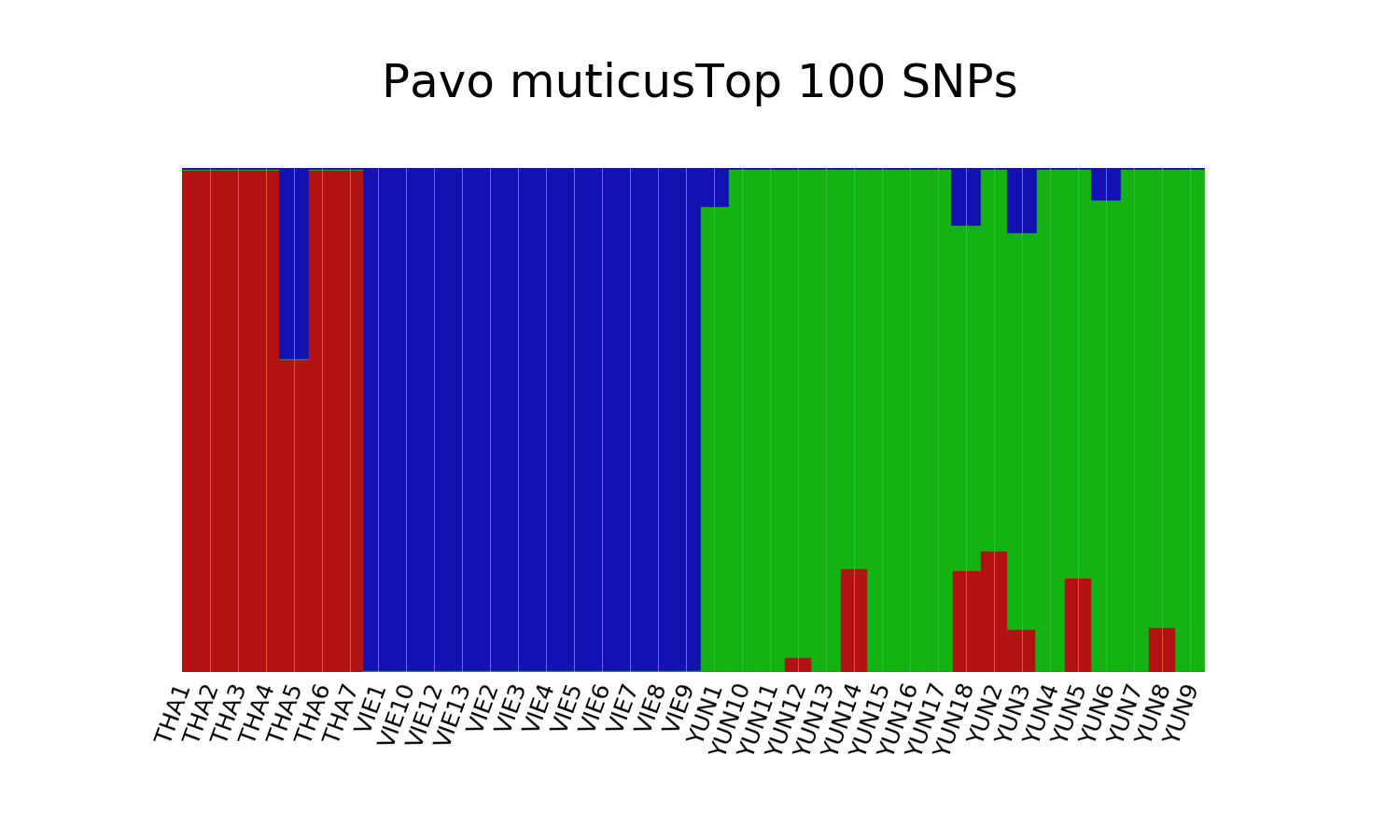

### Supplementary_Figure_2G.png

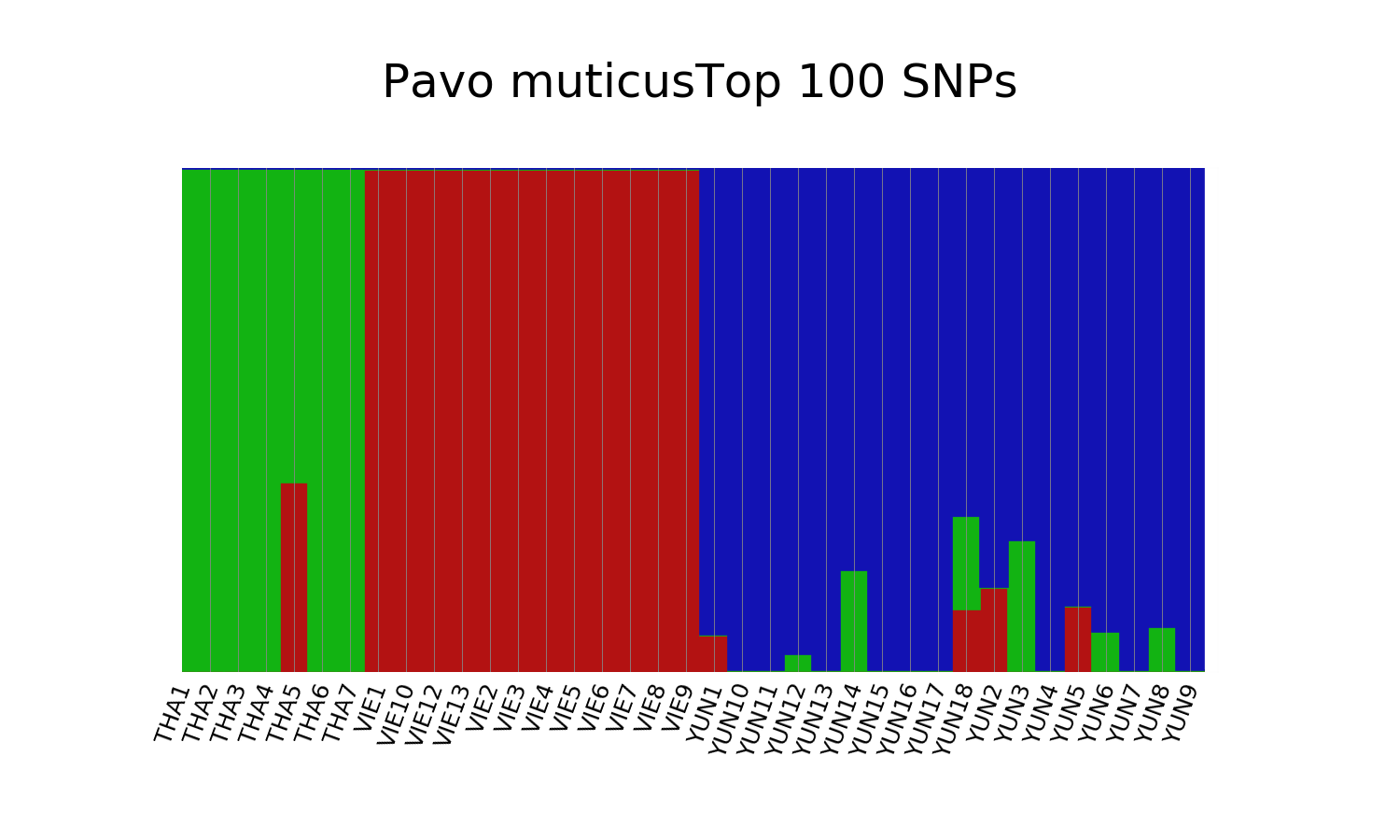

### Supplementary_Figure_2I.png

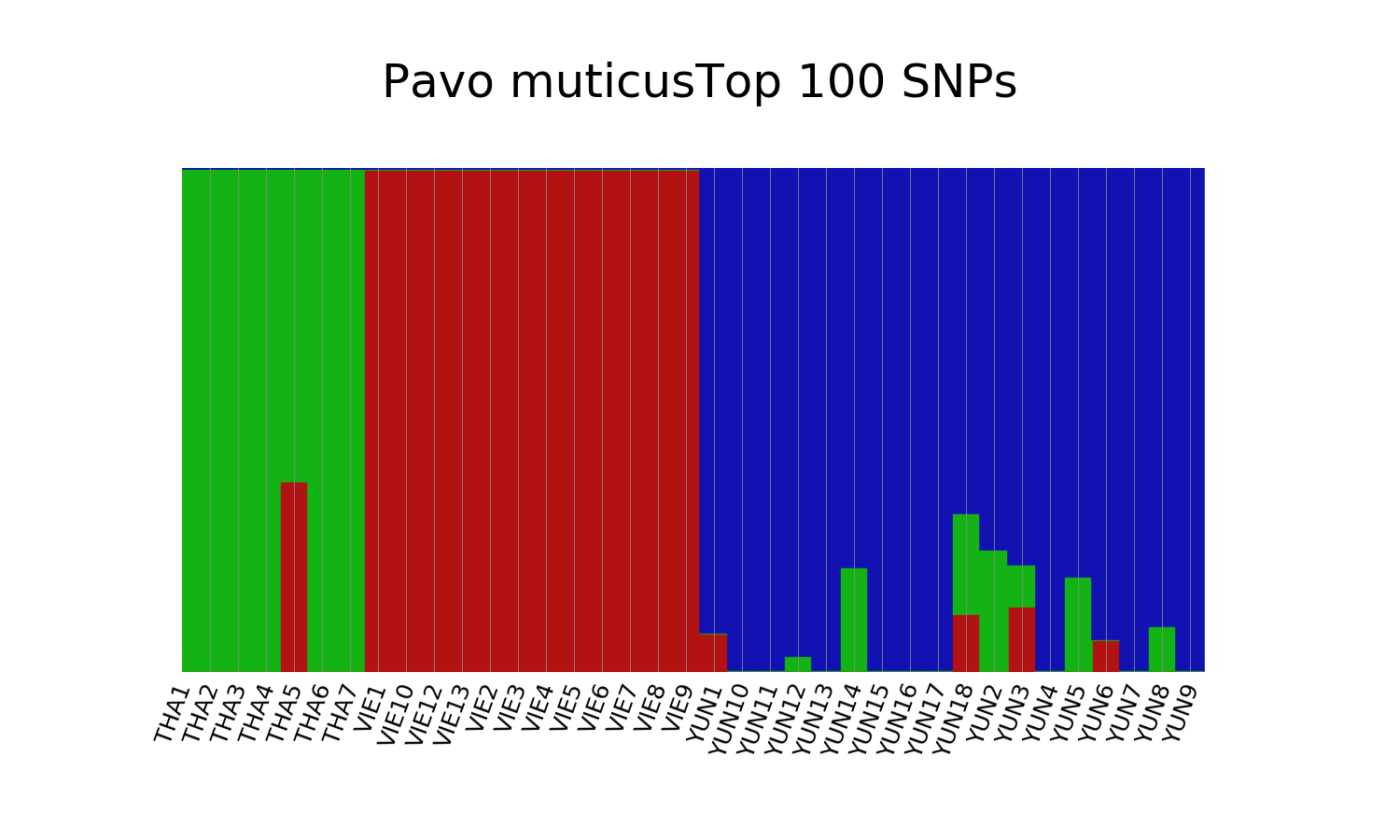

### Supplementary_Figure_3.jpeg

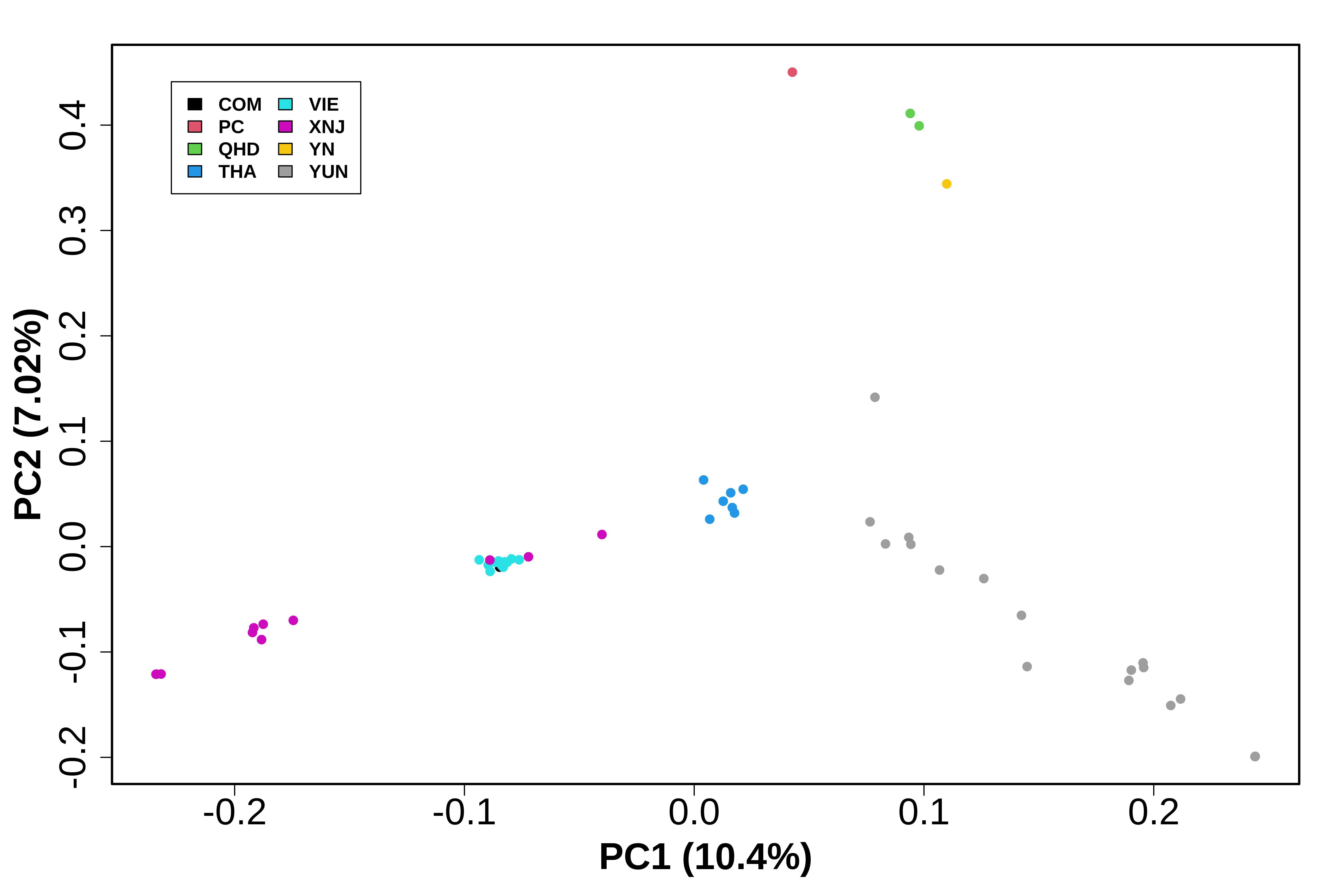

### Supplementary_Figure_4A.png

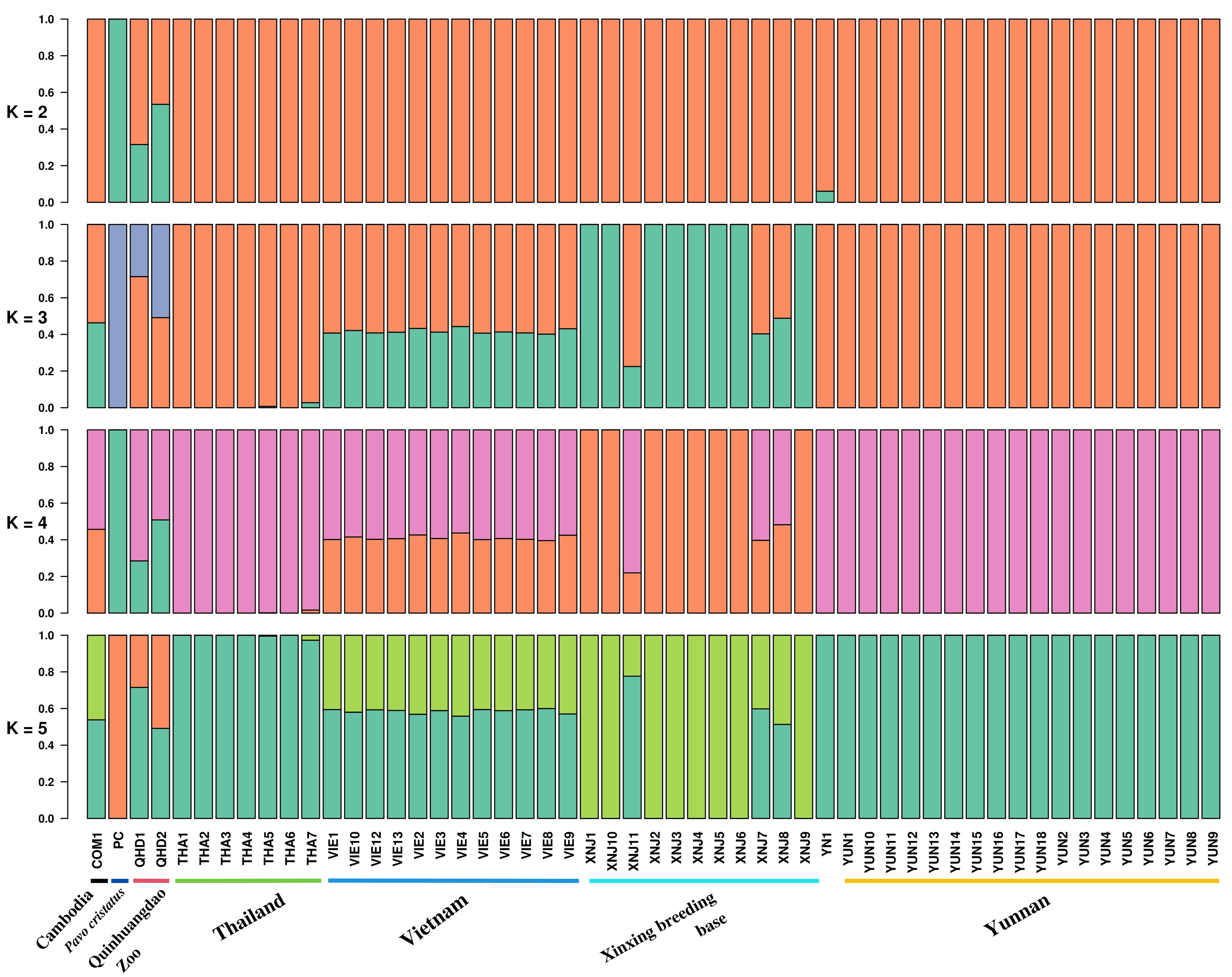

### Supplementary_Figure_4B.png

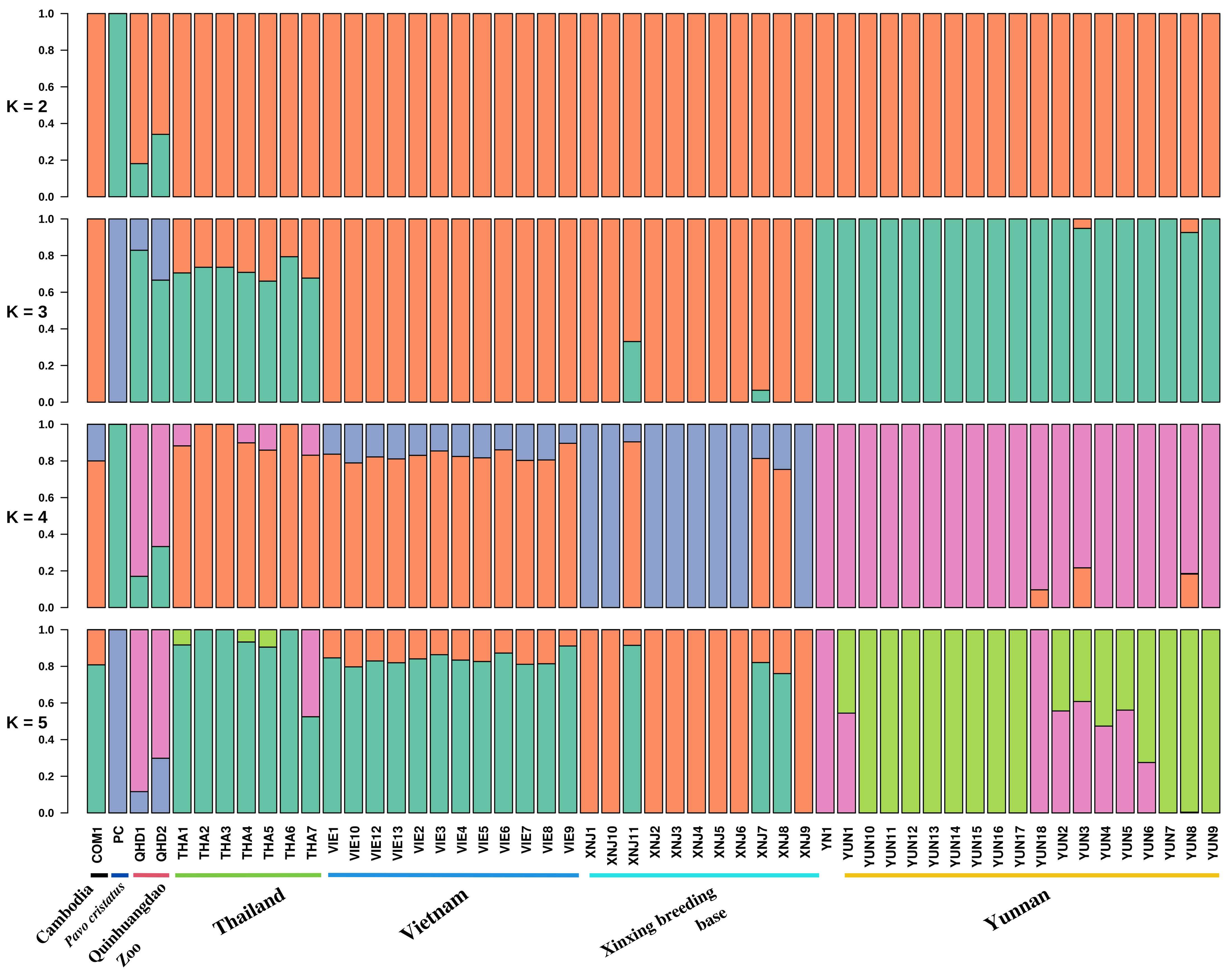

### Supplementary_Figure_5.pdf

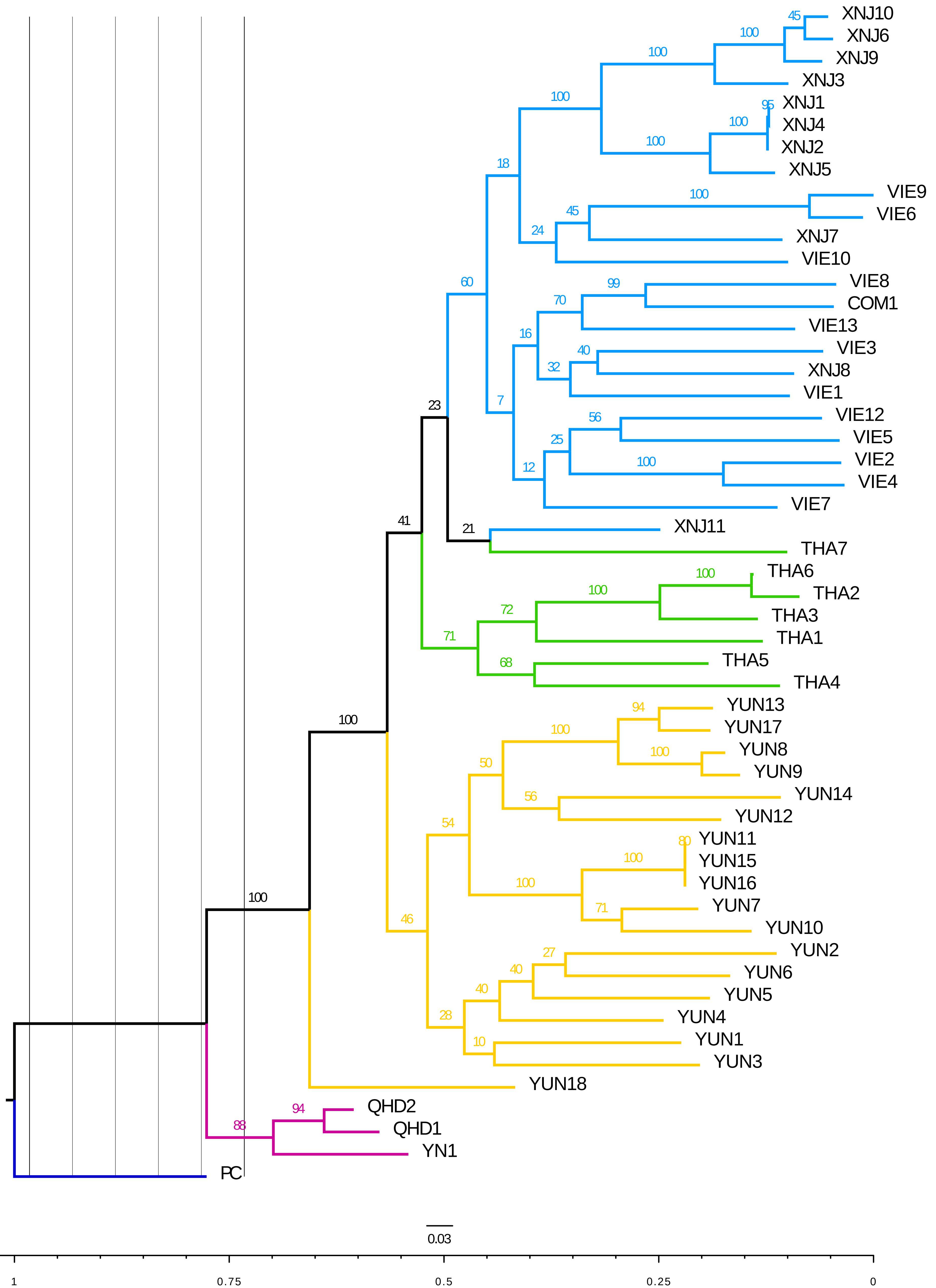

### Supplementary_Figure_6A.jpeg

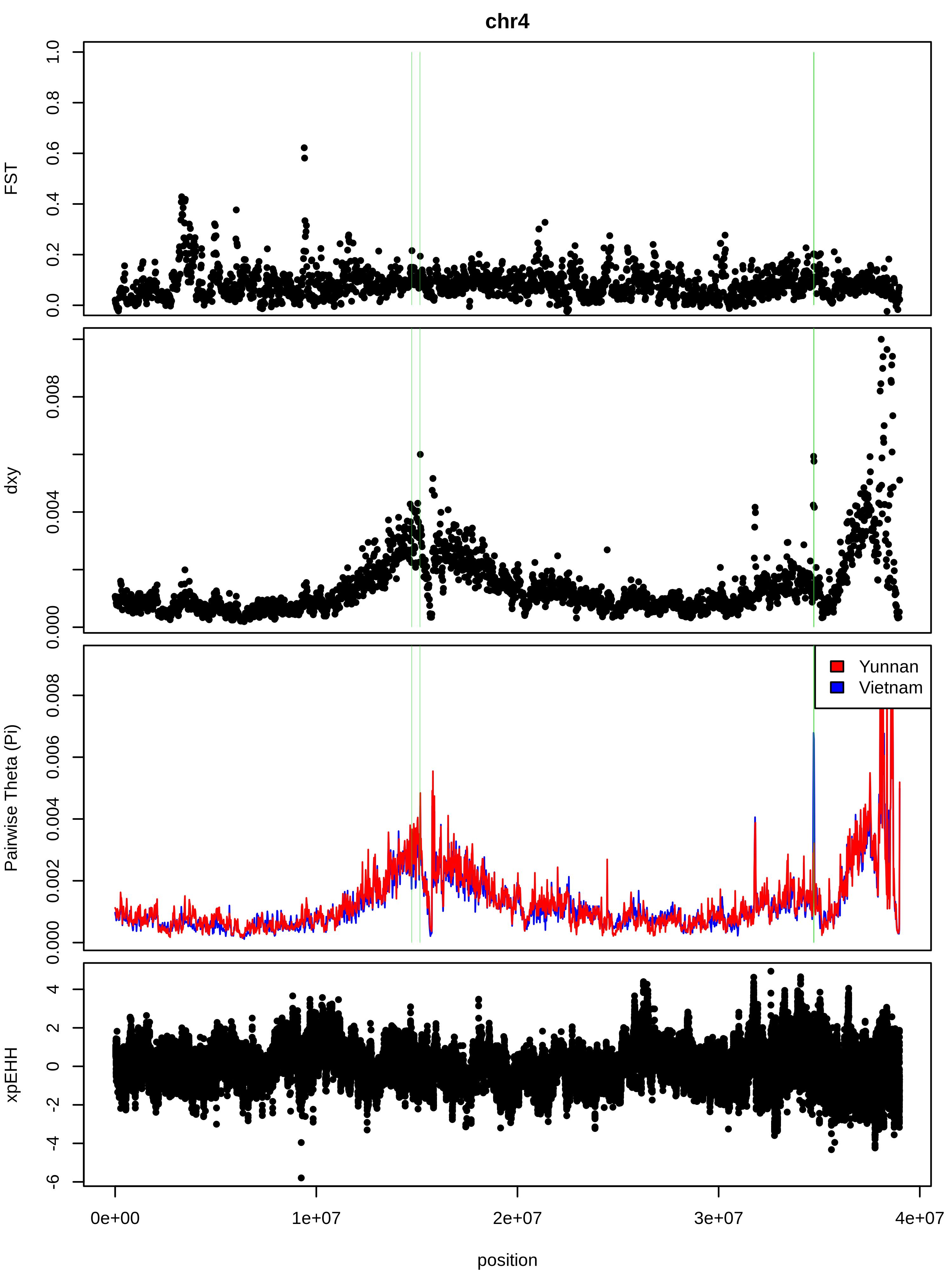

### Supplementary_Figure_6B1.jpeg

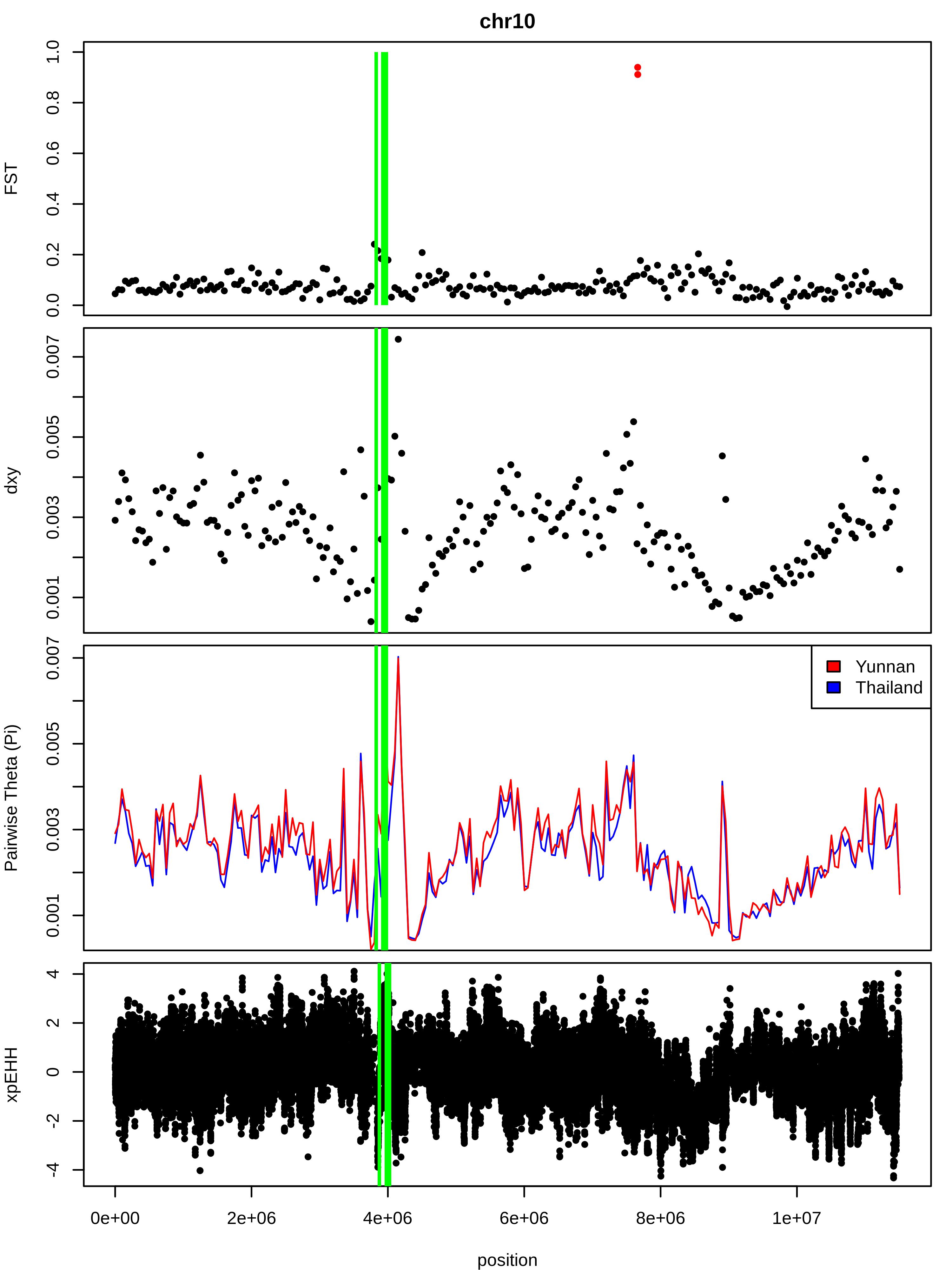

### Supplementary_Figure_6B2.jpeg

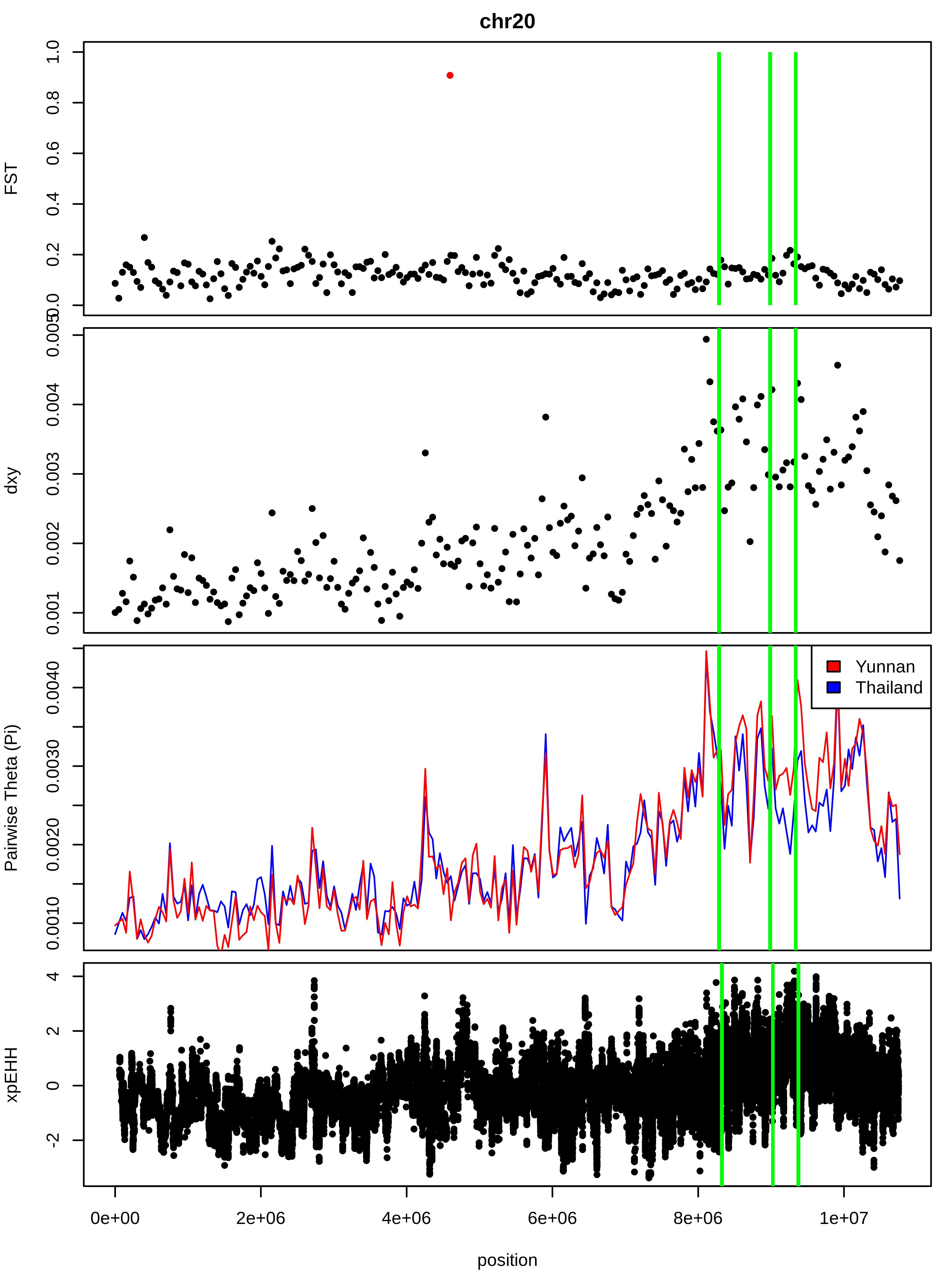

### Supplementary_Figure_6B3.jpeg

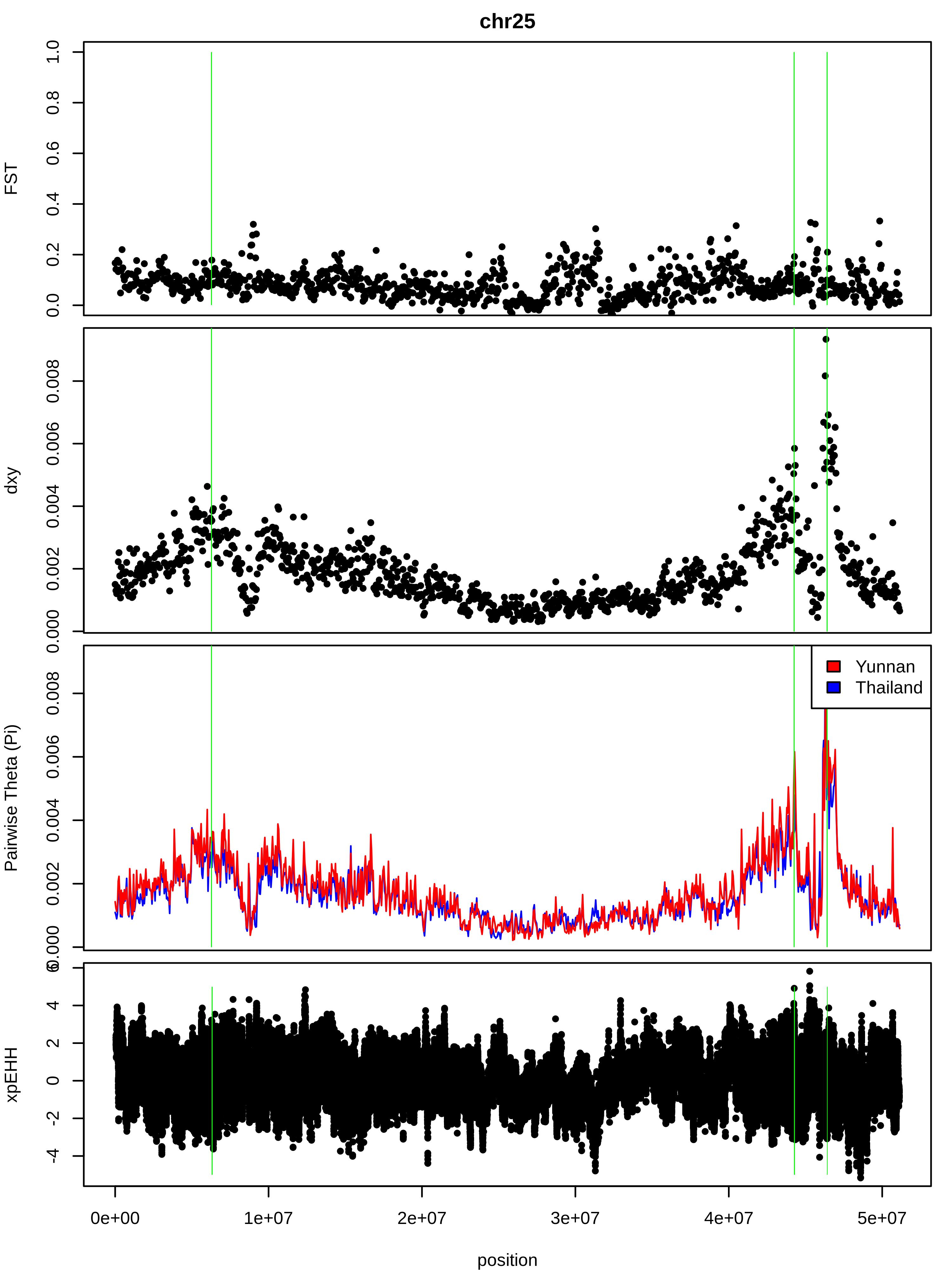

### Supplementary_Figure_6C1.jpeg

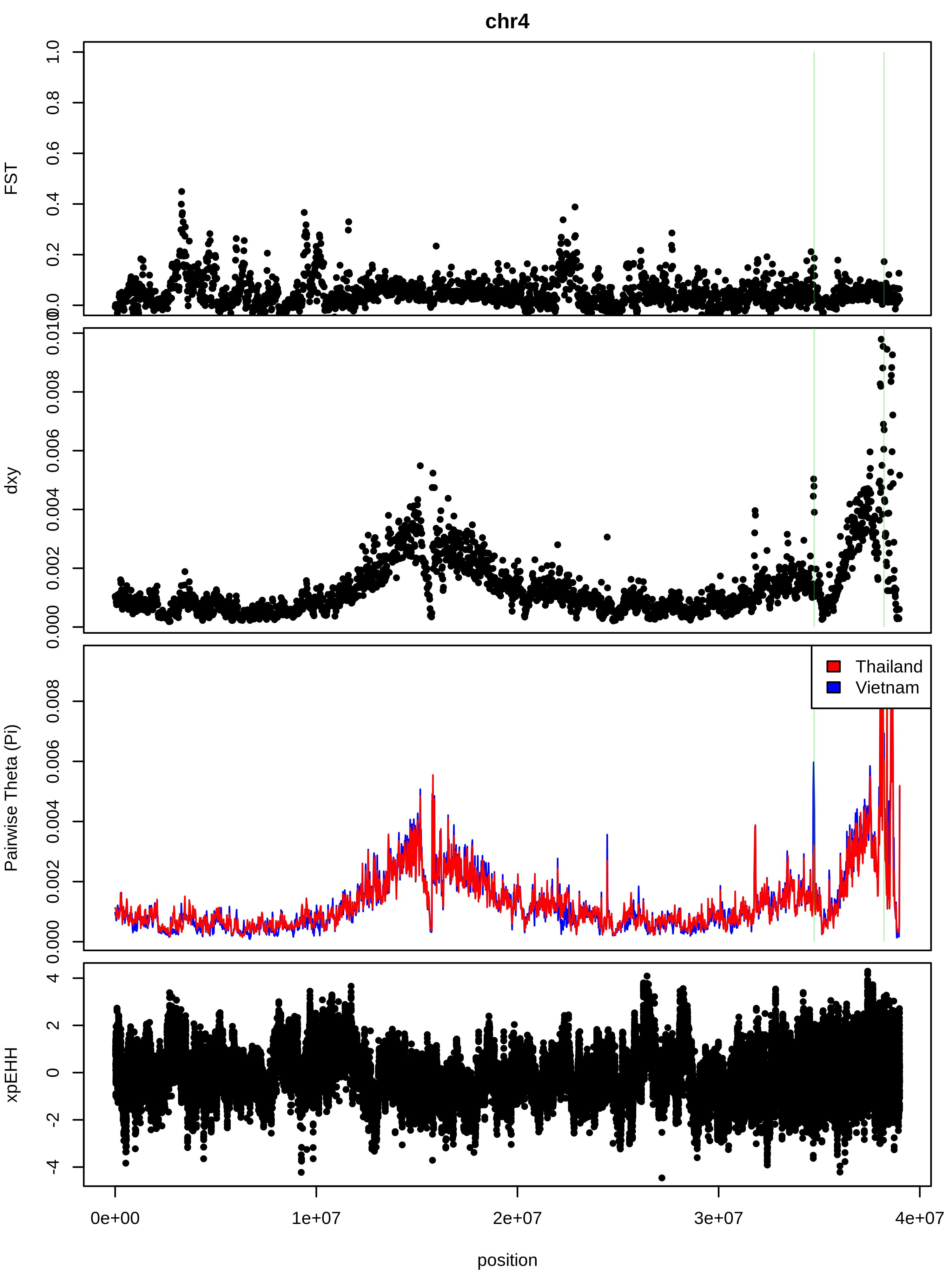

### Supplementary_Figure_6C2.jpeg

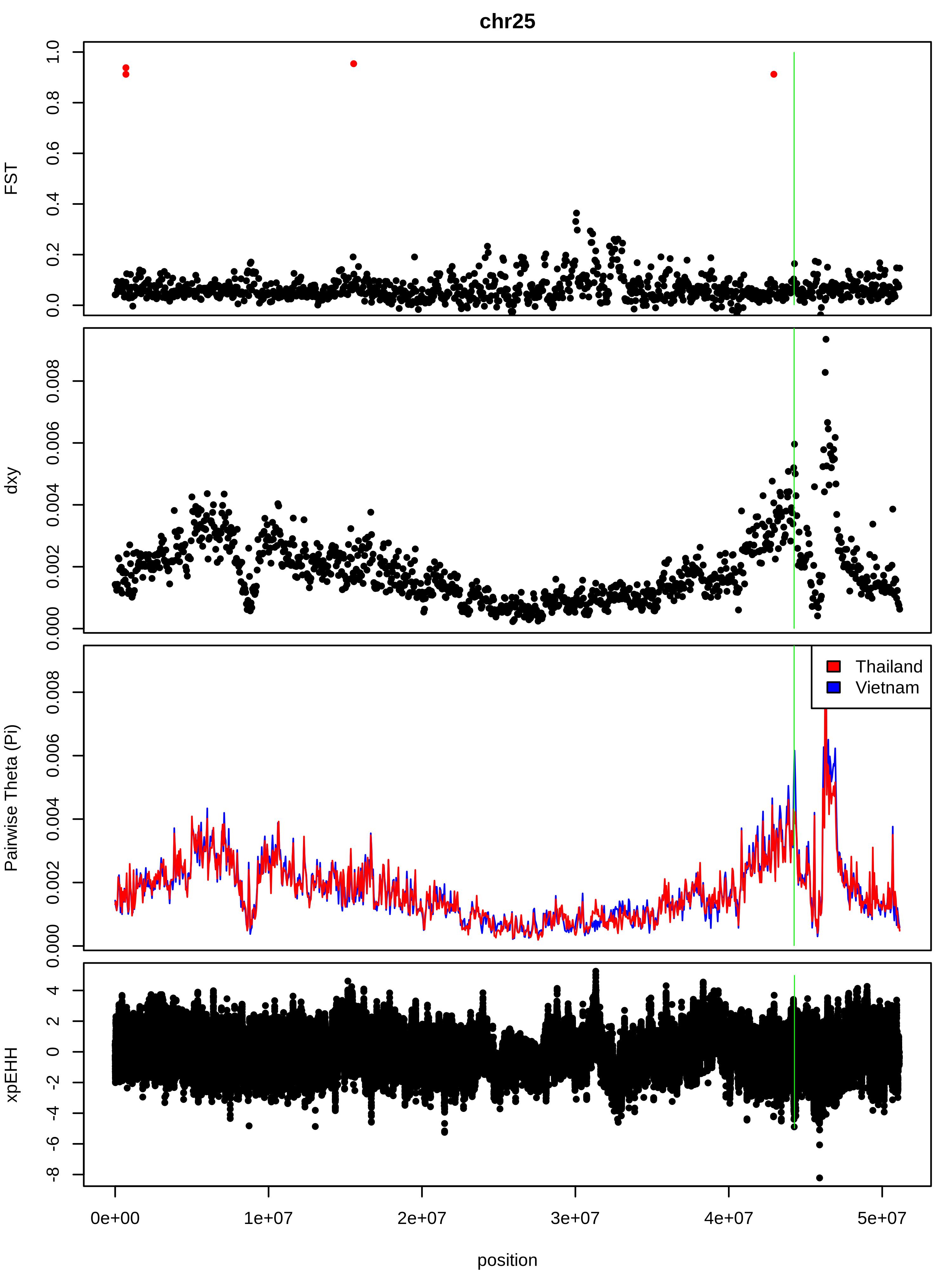

### Supplementary_Figure_7.jpeg

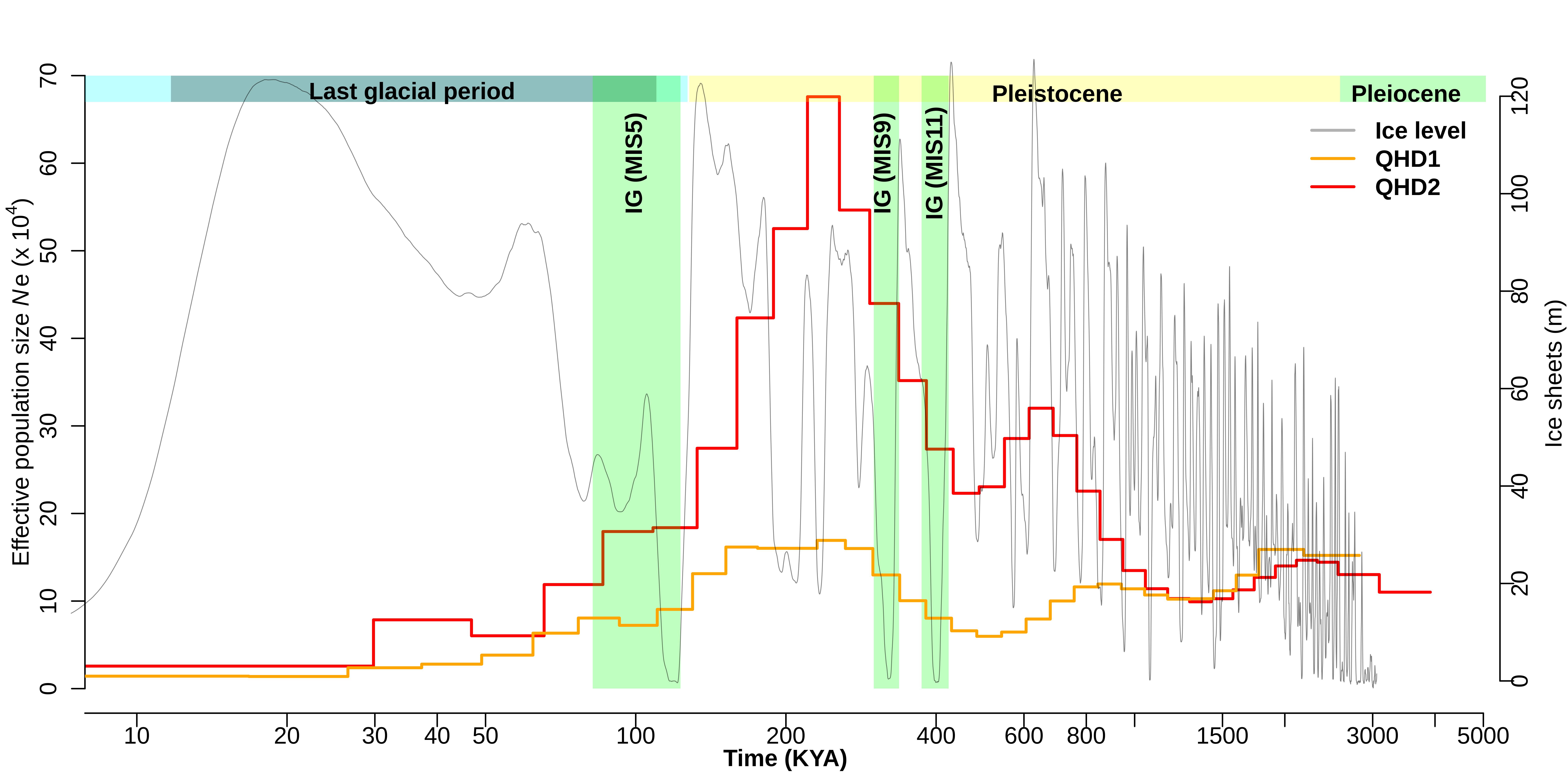

### Supplementary_Figure_8.jpeg

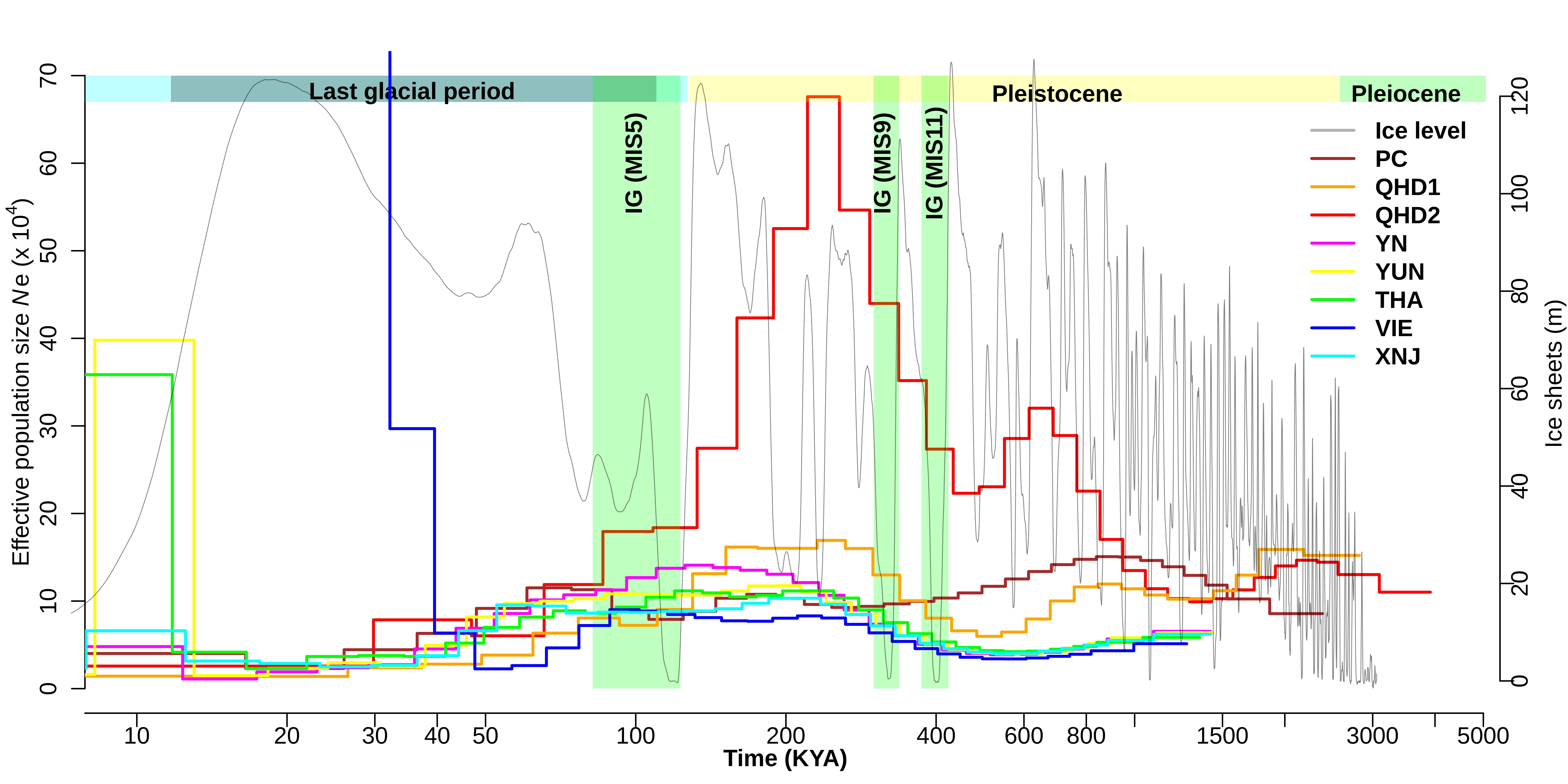

### Supplementary_Figure_9.jpeg

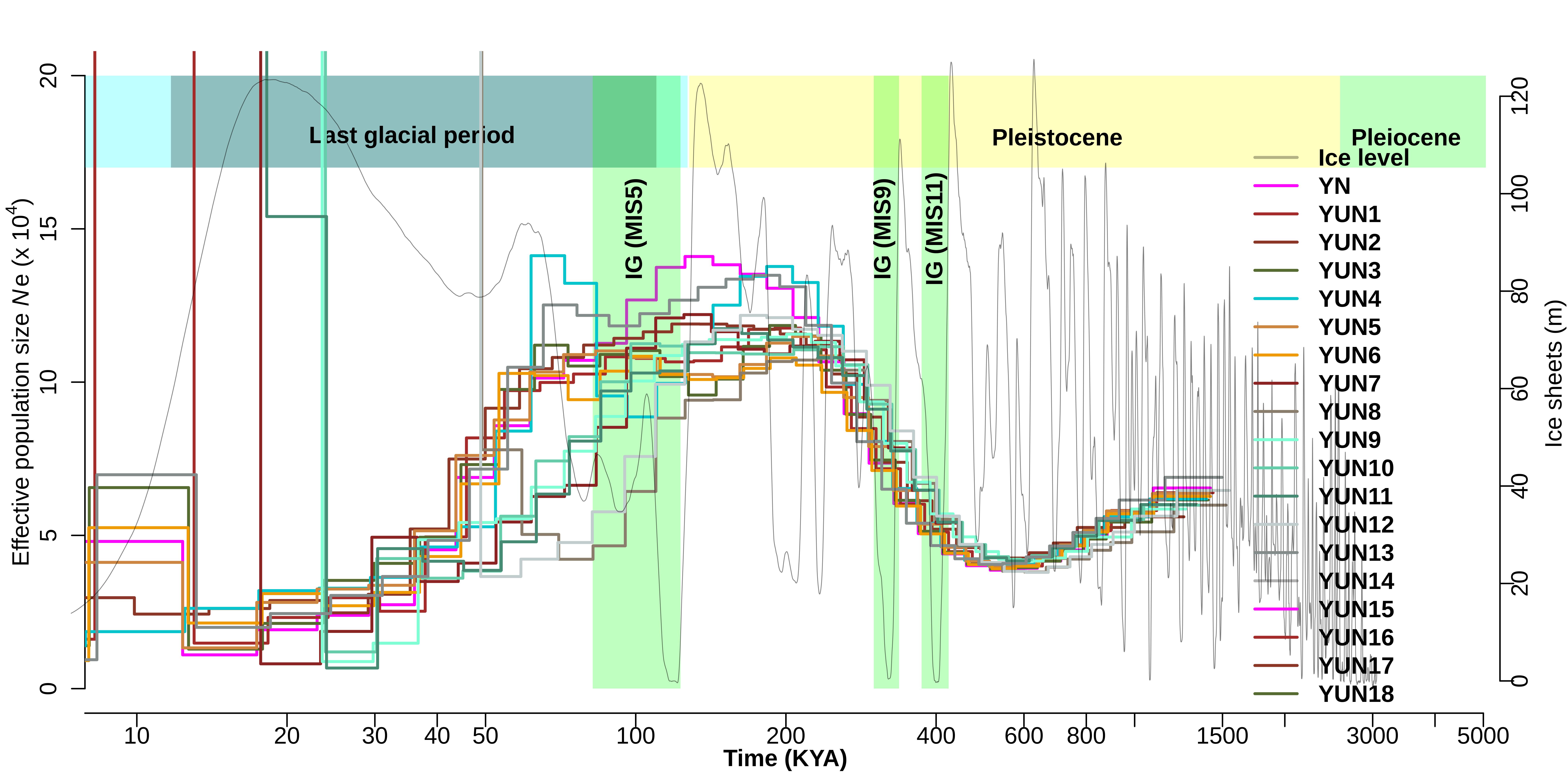

### Supplementary_Figure_10.jpeg

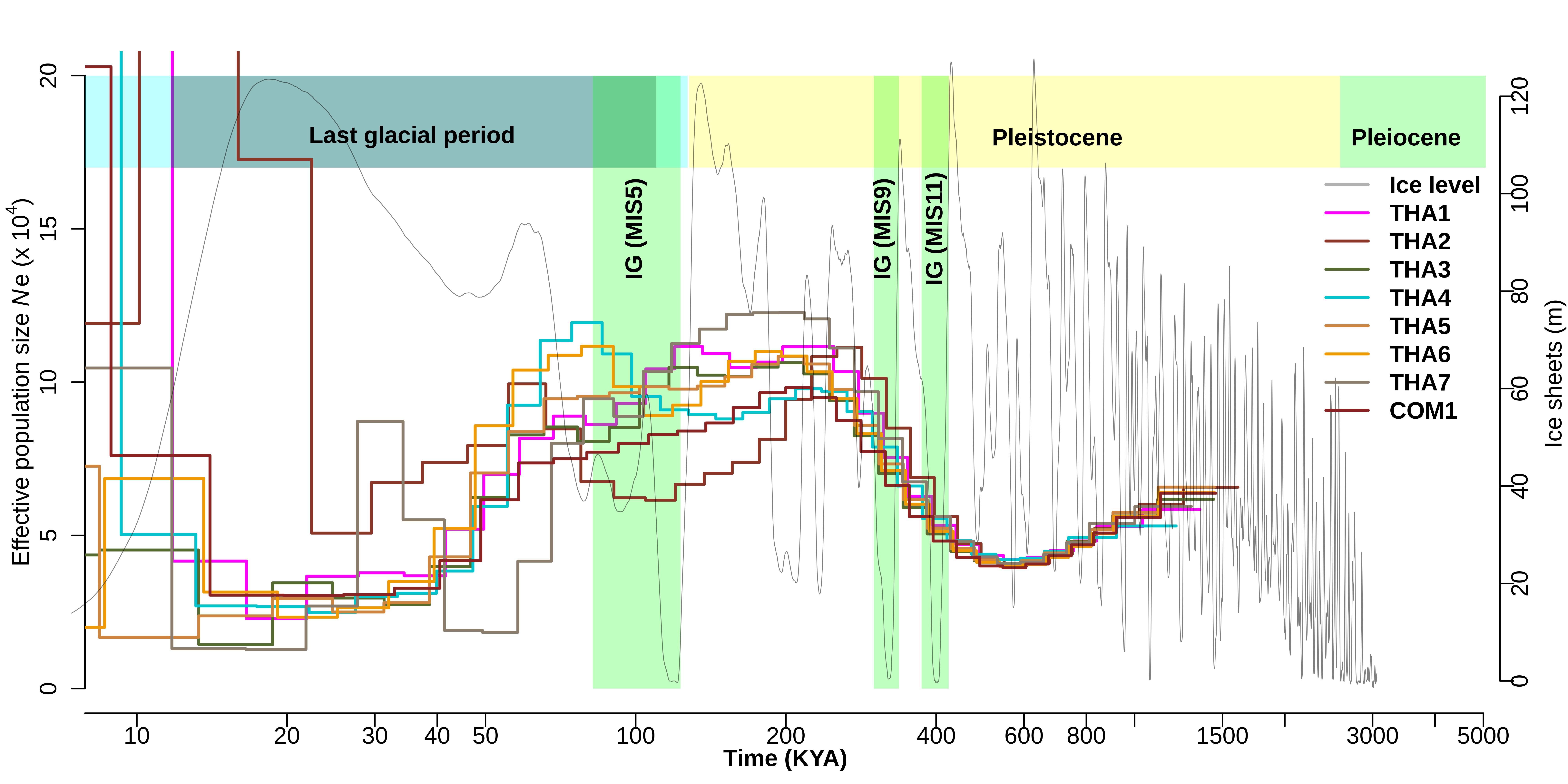

### Supplementary_Figure_11.jpeg

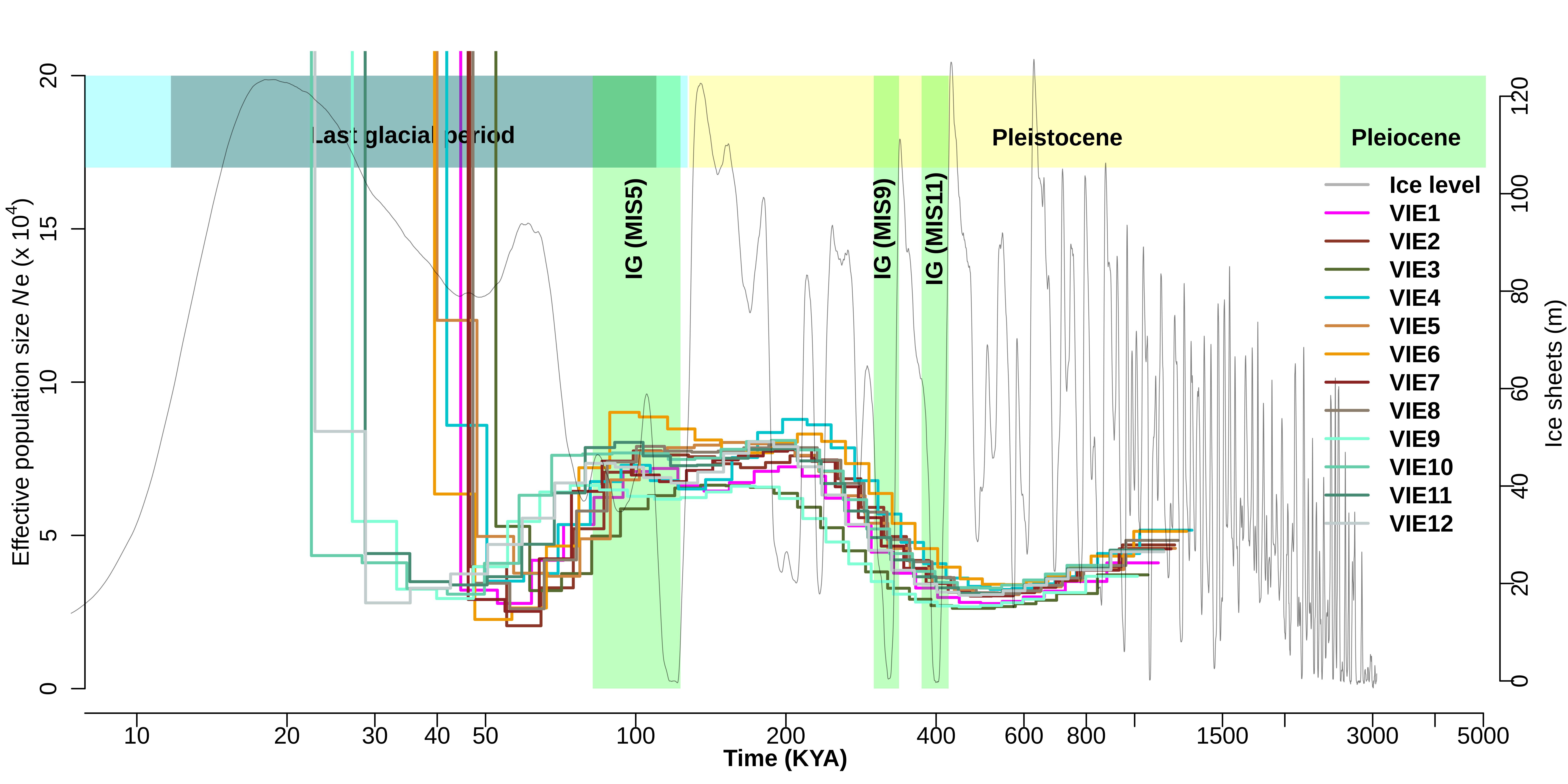

### Supplementary_Figure_12.jpeg

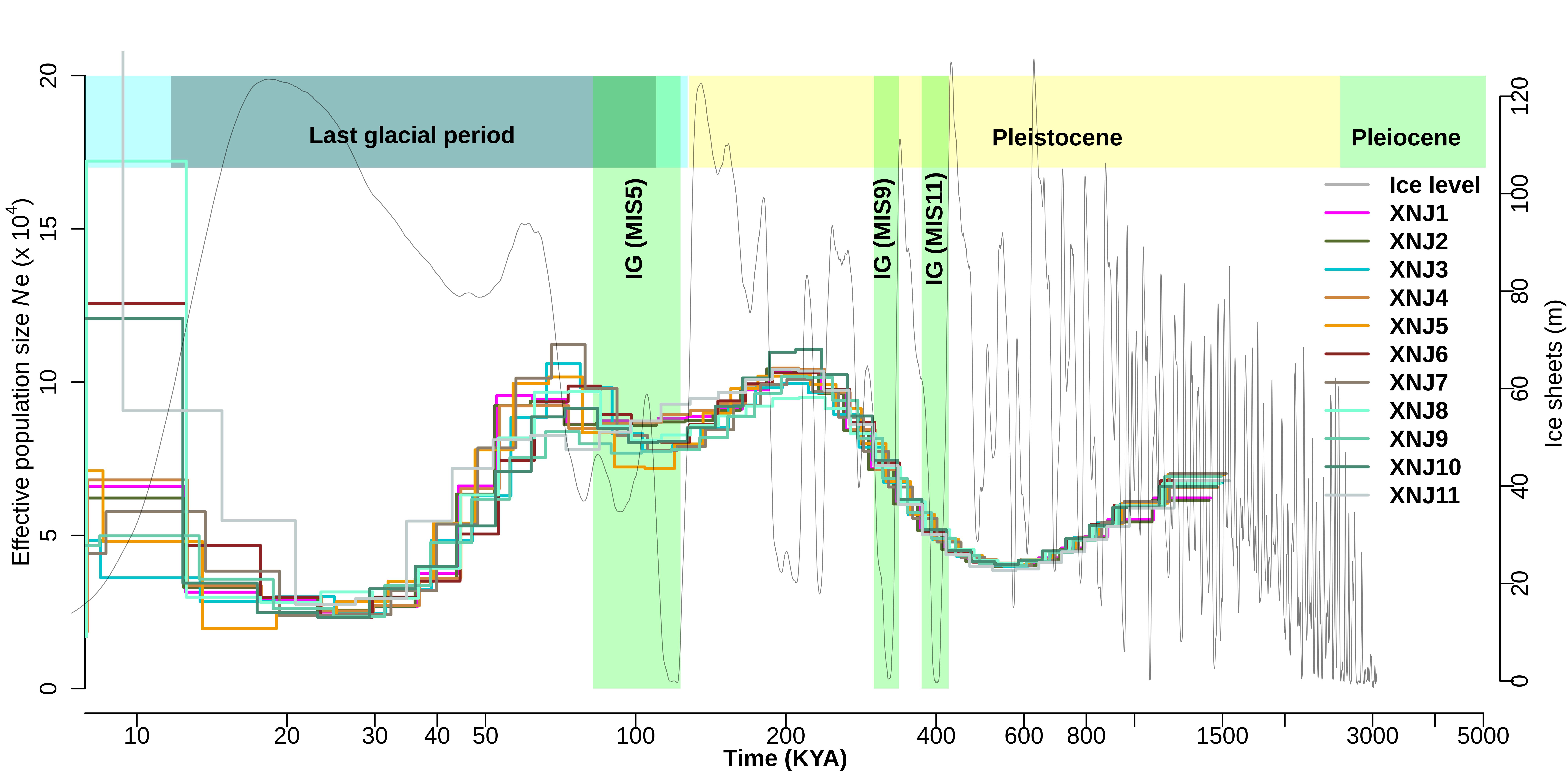

### Supplementary_Figure_13.jpeg

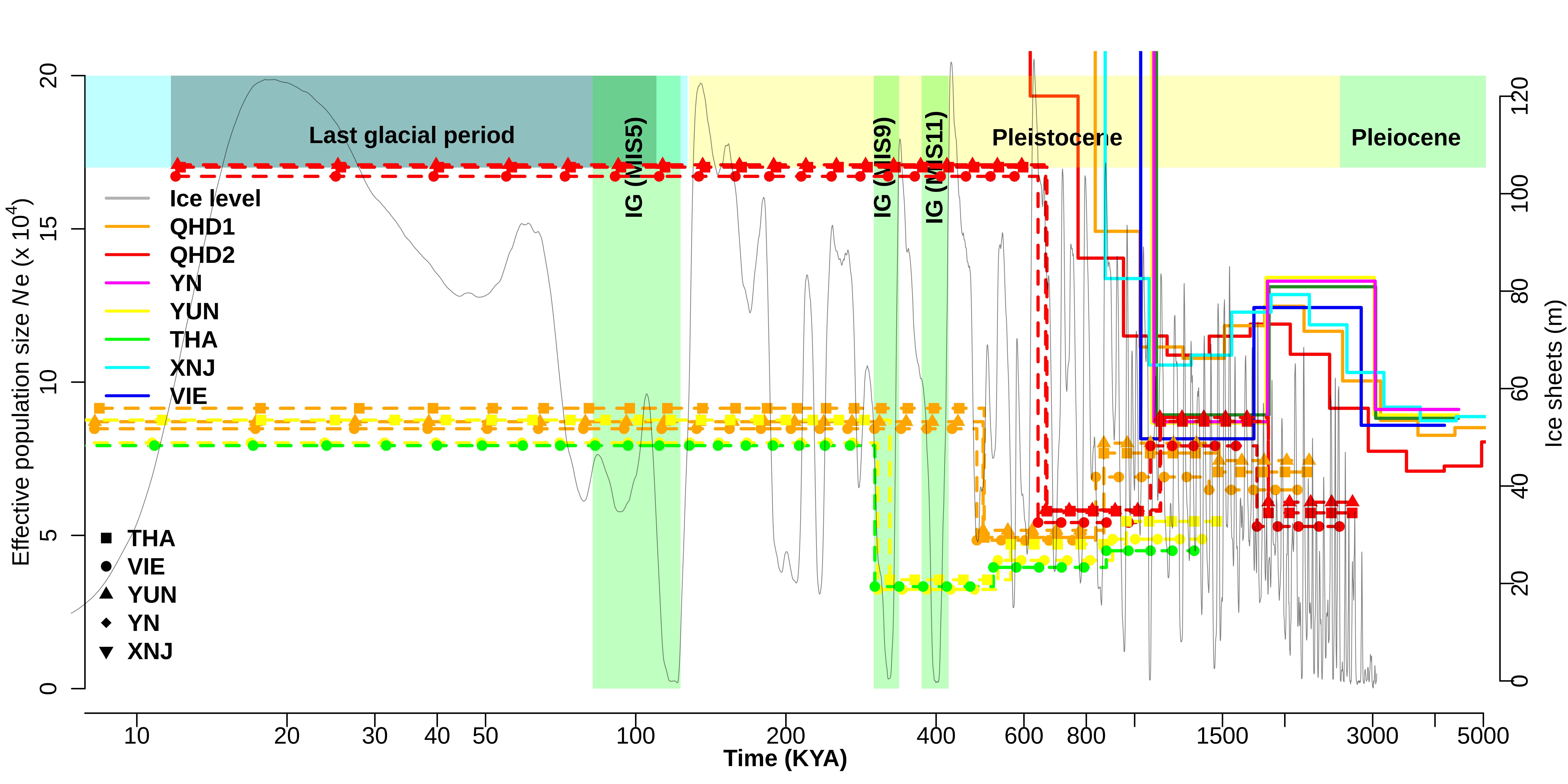

### Supplementary_Figure_14.png

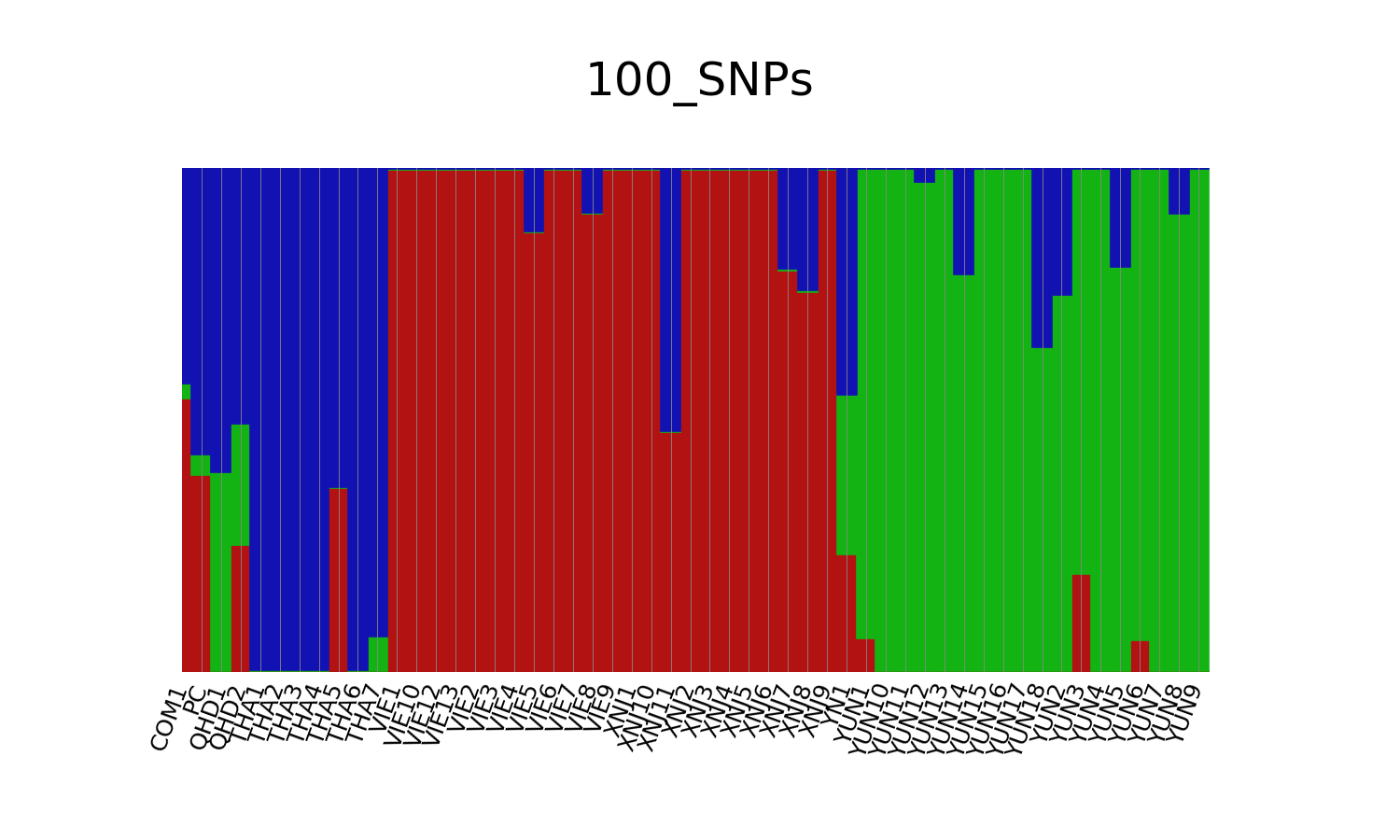

### Supplementary_Figure_15.jpeg

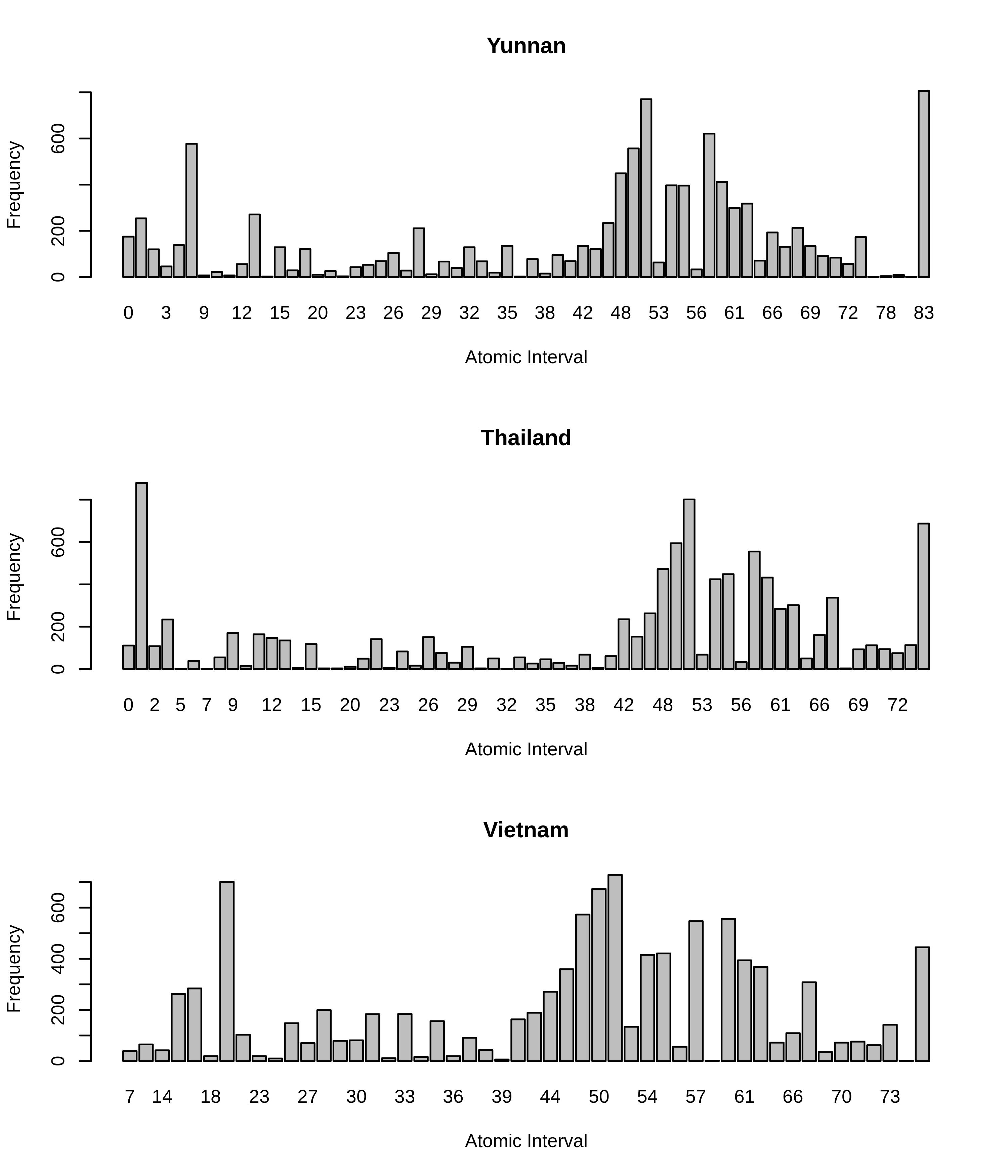
